# Pathogenic DRP1 variants reveal a role for biomolecular condensation in mitochondrial fission

**DOI:** 10.64898/2026.07.06.735726

**Authors:** Kyle A. Ross, Amanda M. Travis, Megan C. Harwig, Micaela S. Young, Erica H. Rodas Montejo, Matthew J. Farah, Michael J. Donohue, Robert W. Taylor, Monika Oláhová, R. Blake Hill

**Affiliations:** Skaggs School of Pharmacy and Pharmaceutical Sciences, University of Colorado Anschutz Medical Campus, Aurora, CO, USA; Department of Cell Biology, Neurobiology, and Anatomy, Medical College of Wisconsin, Milwaukee, WI, USA; Department of Cell & Developmental Biology, Medical College of Wisconsin, Milwaukee, WI, USA; Mitochondrial Research Group, Translational and Clinical Research Institute, Faculty of Medical Sciences, Newcastle University, Newcastle upon Tyne, NE2 4HH, UK; NHS Highly Specialised Service for Rare Mitochondrial Disorders, Newcastle upon Tyne Hospitals NHS Foundation Trust, Newcastle upon Tyne, NE1 4LP, UK; School of Geography and Natural Sciences, Faculty of Science and Environment, Northumbria University, Newcastle upon Tyne, UK

**Keywords:** DRP1, EMPF1, mitochondrial fission, biomolecular condensation, phase separation

## Abstract

Mitochondrial fission is essential for proper organelle function and for cellular homeostasis. Excessive fission is associated with several neurological disorders, whereas impaired fission leads to the lethal encephalopathy EMPF1, which is caused by heterozygous mutations in *DNM1L. DNM1L* encodes the mitochondrial fission mechanoenzyme DRP1, which self-assembles to drive membrane scission. Wild-type DRP1 appear as punctate structures thought to be pre-scission complexes of well-ordered oligomeric assemblies. Immunofluorescence imaging of patient-derived EMPF1 fibroblasts carrying assembly-deficient *DNM1L* variants reveals elongated mitochondrial networks consistent with impaired fission. Despite this loss-of-function phenotype, these cells unexpectedly retain wild-type numbers of DRP1 puncta. We confirmed the previously reported inability of purified pathogenic DRP1 variants p.Gly363Asp and p.Gly401Ser to assemble under conditions in which WT DRP1 forms helical polymers. Under macromolecular crowding conditions, however, both wild-type and mutant DRP1 access condensed states whose formation depends on protein concentration and solution conditions. Acute treatment of EMPF1 fibroblasts with 1,6-hexanediol decreases mitochondrial DRP1 puncta intensity in mutant cells relative to wild type, indicating genotype-dependent differences in puncta. The magnitude of these differences supports a dominant-negative phenotype of the pathogenic variants. Together, these findings support a model in which DRP1 puncta occupy a continuum of states, from liquid-like condensates to ordered, fission-competent assemblies, revealing biomolecular condensation as a previously unrecognized layer of DRP1 regulation. Biasing DRP1 along this continuum may provide a mechanistic basis for impaired fission in EMPF1 and suggest opportunities to restore productive assembly in select pathogenic contexts.

**Significance Statement:** DRP1 puncta associated with mitochondrial fission are thought to be well-ordered oligomeric assemblies that precede membrane scission. Yet their dynamic behavior within cells has remained difficult to reconcile as well-ordered assembly. Under prevailing models, cells bearing pathogenic *DNM1L* variants impaired in assembly would be expected to lack puncta, but we show these cells retain wild-type puncta levels. We demonstrate that both wild-type and pathogenic mutant DRP1 populate multiple condensed states *in vitro*, and that disease variants are biased toward more fluid, chemically sensitive assemblies. These findings identify biomolecular condensation as a regulatory layer of DRP1 organization and suggest that shifting DRP1 along this assembly continuum may restore productive fission in select pathogenic contexts.

## Introduction

Mitochondrial fission is essential for organellar function and eukaryotic cell homeostasis through several pathways, such as apoptosis, mitophagy, immune signaling, and cell cycle regulation (1–3). Increased mitochondrial fission is linked to Alzheimer’s, Parkinson’s, and Huntington’s disease, and to other neurological disorders (4, 5). By contrast, decreased mitochondrial fission from heterozygous mutations in the mitochondrial fission gene *DNM1L* cause EMPF1 (<u>e</u>ncephalopathy due to defective <u>m</u>itochondrial and <u>p</u>eroxisomal fission-<u>1</u>, MIM #614388) and results in seizures, hypotonia, and severe neurological decline that often leads to childhood mortality (6, 7). *DNM1L* encodes <u>d</u>ynamin-related <u>p</u>rotein <u>1</u> (DRP1, or Dnm1 in yeast), which is a member of the dynamin superfamily and the GTPase mechanoenzyme that mediates mitochondrial and peroxisomal fission (8–12). Consistent with an essential role for DRP1, *Dnm1l* loss is lethal in mice and disrupts brain development (13), and disease-associated *DNM1L* variants produce elongated mitochondrial networks consistent with impaired fission (14–18).

Endogenous DRP1 in cells visualized by immunofluorescence microscopy is diffusely distributed throughout the cytoplasm and is concentrated in discrete puncta, or foci, on mitochondria (15, 19–22) and peroxisomes (10–12, 17). GFP-tagged DRP1 similarly accumulates into punctate foci at mitochondrial division sites, where DRP1 is required for mitochondrial fission (10, 23). Mitochondrial metabolic state and stress influence DRP1 puncta distribution and brightness (24, 25). In yeast, Dnm1-GFP forms bright, cortical puncta that colocalize with mitochondrial tubules and appear at fission sites (8, 9, 26). Live-cell movies showed that Dnm1 puncta assemble rapidly, reside transiently at fission sites, and dissipate after scission (27). The physical basis of puncta formation is thought to derive from the well-characterized ability of DRP1 to self-assemble, similar to other soluble dynamin family members (28–30). Crystal structures of DRP1 defined oligomerization interfaces compatible with dynamin-like oligomerization (31–34). Cryo-electron microscopy further revealed the formation of helical spirals, rings, and membrane-bound assemblies that encircle and constrict membranes to defined diameters (35–40). Together, these biochemical and biophysical studies established that DRP1 can form well-ordered oligomeric assemblies capable of membrane constriction, and motivated the view that mitochondria-associated DRP1 puncta correspond to fission-competent pre-scission complexes.

However, a model in which all DRP1 puncta are comprised of rigid helical high-order oligomers is not consistent with multiple live-cell studies establishing that DRP1 puncta are heterogeneous (20, 21). Fluorescence recovery after photobleaching (FRAP) studies showed that GFP-DRP1 puncta exchange subunits rapidly with the cytoplasmic pool and that perturbations in the intrinsically disordered variable domain of DRP1 alter puncta dynamics (20, 21, 41). Single-particle tracking and live-cell imaging revealed that DRP1 monomers within puncta exhibit rapid lateral mobility along mitochondrial tubules, inconsistent with static higher-order assemblies (21). This work showed that DRP1 matures progressively on mitochondria from small, mobile assemblies that merge into larger, stable oligomers, only a small subset of which catalyzes membrane scission. Super-resolution, live-cell imaging further identified two functionally distinct classes of DRP1-mediated fission: Peripheral fission that segregates damaged mitochondria for degradation and midzone fission that promotes mitochondrial biogenesis (25). These findings demonstrate that DRP1 puncta are functionally heterogeneous. More generally, numerous DRP1 puncta occur in the cytoplasm, not on mitochondria or peroxisomes, which is difficult to reconcile with a purely pre-scission role. Fluorescence correlation spectroscopy supports the presence of low-order cytoplasmic DRP1 complexes (e.g., dimers/tetramers) (42, 43), but how such pools relate to the formation of bright cytoplasmic puncta remains unclear. These studies report a variety of DRP1 properties that are difficult to reconcile with a single static assembly state in puncta, but elevated expression levels or the presence of non-native fusion tags may perturb DRP1 organization.

We and others have undertaken analyses of native DRP1 at endogenous levels in both healthy and diseased cells by immunofluorescence to probe how native puncta form and behave. Patient-derived dermal fibroblast cells harboring the heterozygous *DNM1L* (NM_012062.5) variants c.1088G>A, p.Gly363Asp (G363D) or c.1201G>A, p.Gly401Ser (G401S) show elongated mitochondrial and peroxisomes with modest differences in DRP1 puncta distributions (17). In a different study, examination of the activating phosphorylation of DRP1 at Ser616 revealed that the quantity and volume of mitochondrially-associated pS616-DRP1 puncta are not markedly altered in cells carrying pathogenic *DNM1L* variants c.95G>C, p.Gly32Ala (G32A) or c.1207C>T, p.Arg403Cys (R403C) (18). Given that the variants G363D, G401S, and R403C are assembly-deficient *in vitro* (15, 17, 44), the presence of apparently normal subcellular puncta is particularly striking. These findings challenge the classical view that DRP1 puncta arise solely from the well-ordered helical assembly at fission sites and underscore the need to better define the physical basis of puncta formation in cells.

Here, we examine DRP1 puncta in patient-derived EMPF1 fibroblasts carrying the G363D or G401S alleles, and quantify DRP1 puncta number, intensity, and spatial distribution at endogenous expression levels. We demonstrate that the cells carrying assembly-deficient DRP1 variants form essentially wild-type numbers of puncta. We examined purified DRP1 constructs under a variety of conditions seeking a physical basis for puncta formation. We confirmed the inability of the mutants to assemble under standard conditions, identified a latent capacity for G401S (but not G363D) to self-assemble at low ionic strength, and found that the wild-type and mutant proteins can form condensed phases. Pathogenic DRP1 variants enter these condensed states more readily than wild-type, raising the possibility that condensation contributes to DRP1 participation in puncta. We show that 1,6-hexanediol, which is known to perturb biomolecular condensates in cells, strongly perturbs DRP1 subcellular distribution in cells carrying assembly-deficient DRP1 variants and modestly perturbs DRP1 subcellular distribution in wild-type fibroblasts, although a more pronounced effect is seen in other cell types at endogenous levels. Together, these observations support an extension of the current model in which DRP1 puncta derive, at least in part, from a condensed phase, potentially a biomolecular condensate similar to other systems (45, 46). That pathogenic variants appear to occupy these condensed states more readily than wild-type suggesting a maturation-trap model for their dysfunction and hints at strategies that might restore more normal assembly behavior.

## Results

To establish a quantitative framework for analyzing DRP1 puncta, we first revisited the cellular phenotypes of pathogenic *DNM1L* variants using high-resolution confocal microscopy (**Fig. 1A**). As reported previously, fibroblasts carrying the G363D and G401S variants exhibited elongated mitochondrial networks relative to healthy controls, which was not due to reduced total DRP1 levels as assessed by Western blot analyses (17). Quantitative analysis using MitoGraph (47) confirmed a significant increase in average mitochondrial length from 6.9 ± 2.9 µm to 9.7 ± 6.3 µm and 12.4 ± 6.8 μm in G363D and G401S cells, respectively (**Figs. 1B*i* and S1**). Although overall DRP1 immunofluorescence showed reduced colocalization with mitochondria in patient cells (**Figs. S1, S2**), we reasoned that global colocalization metrics provide limited insight into how DRP1 organizes into discrete fission-related structures, given the extensive mitochondrial volume and the largely cytoplasmic distribution of DRP1. We therefore focused on DRP1 puncta, which are widely interpreted as functionally relevant assemblies. Immunostaining for DRP1 revealed an unexpectedly large number of endogenous puncta, averaging ∼1,571 in healthy control cells (**Fig. 1B*ii***). Puncta numbers were unchanged in G363D fibroblasts, and only modestly reduced in G401S cells (∼1,303 puncta), despite both variants being defective in ordered, self-assembly *in vitro* (15, 17, 48). To determine whether puncta distribution was altered, we implemented autonomous three-dimensional puncta detection. TOM20 immunofluorescence was used to generate 3D mitochondrial masks (and their inversions), which were combined with the DRP1 immunofluorescence signal to distinguish mitochondrial-associated from cytoplasmic puncta. Surprisingly, only ∼30% of DRP1 puncta were associated with mitochondria in control cells (**Fig. 1B*iii***), indicating that the majority of DRP1 puncta form independently of mitochondrial recruitment. Patient-derived cells showed a modest reduction in mitochondria-associated puncta (to ∼27%), but this difference was minor relative to the overall predominance of non-mitochondrial puncta across all genotypes. DRP1 also mediates peroxisomal fission and has been reported to be enriched at other subcellular structures including the endoplasmic reticulum, lysosome, and microtubules. Immunostaining colocalization analyses for these structures showed modest colocalization of DRP1 (Pearson’s correlation coefficients < 0.3 regardless of genotype, **Fig. S2**), indicating that a majority of puncta are likely cytoplasmic. Notably, mitochondrial-associated DRP1 puncta were consistently brighter (∼1.5-fold or more) than the non-mitochondrial population in both control and patient cells (**Fig. 1B*iv***). This intensity difference suggests that DRP1 puncta associated with mitochondria represent a distinct, higher-density population.

**FIGURE 1.**
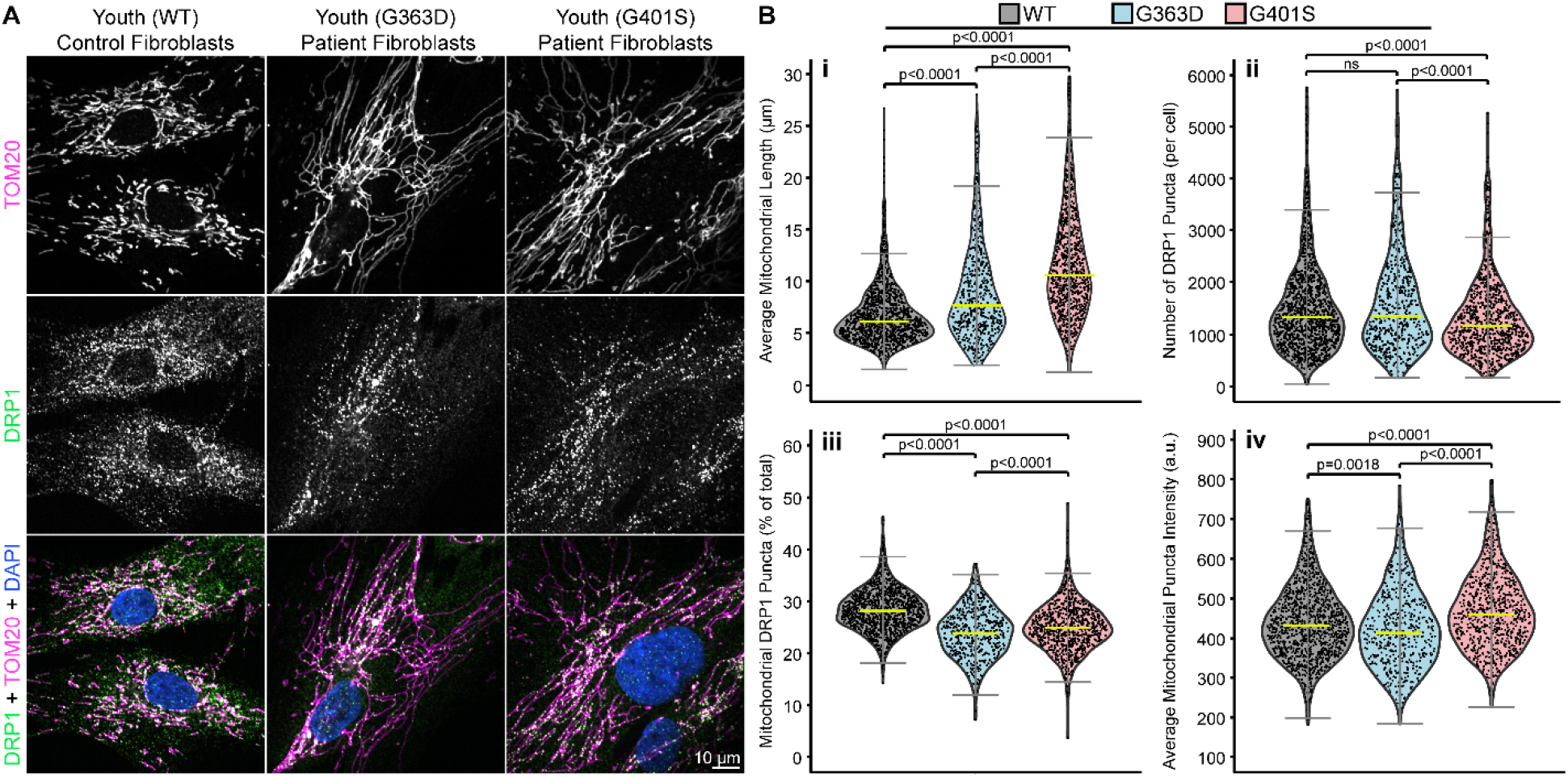
Pathogenic DRP1 variants in patient fibroblasts form abundant endogenous puncta despite mitochondrial elongation. **(A)** Representative fluorescent images of untreated human skin fibroblasts from a healthy age-matched control (wild-type DRP1, left) and from patients carrying the DRP1 G363D (middle) or G401S (right) variants. Cells were immunostained for TOM20 (top row, magenta) and DRP1 (middle row, green), and nuclei were stained with DAPI (blue). Identical LUTs were assigned for each image. Scale bar = 10 µm. **(B)** Quantitative metrics collected from cell images in (A) of untreated human skin fibroblasts for DRP1 WT (gray), G363D (light blue), and G401S (light red). Average mitochondrial length (i) was obtained using MitoGraph software (see methods) (n = 1240, 494, or 833 cells for DRP1 WT, G363D, and G401S, respectively). The total number of DRP1 puncta (ii), percentage of mitochondrial DRP1 puncta (iii), and mitochondrial DRP1 puncta intensity (iv) metrics were all obtained using the 3DSuite plugin in FIJI (see methods) (n = 1237, 494, or 831 cells for DRP1 WT, G363D, and G401S, respectively). All statistical tests were performed using one-way ANOVA with Tukey’s post-hoc multiple comparisons test, and p-values are indicated for each comparison (ns = not significant). Yellow bars represent the population median. Error bars are ± 2SD. Source data for this figure are included in supplemental table **Dataset_S01**.

The discovery that assembly-impaired pathogenic *DNM1L* variants form puncta is striking, considering the prior assumption that puncta represent well-ordered, pre-scission complexes (10, 21, 23, 26, 28–34, 41, 49–51). To investigate this further, we purified recombinant proteins for biochemical and biophysical analyses. We first confirmed prior observations that G363D and G401S are assembly-impaired (15, 17, 48). To assess self-assembly, we added the non-hydrolyzable GTP analog, GMP-PCP, which stimulates assembly (21, 32, 35, 39, 41), and evaluated assembly by sedimentation, with larger complexes present in the pellet (**Fig. 2A**). Without nucleotide, all DRP1 constructs remained predominantly in the soluble supernatant fraction, with minimal pelleting observed. After incubation with GMP-PCP, however, DRP1 WT shifted to a roughly equal distribution between the supernatant and pellet, indicative of nucleotide-stimulated self-assembly. By contrast, the pathogenic DRP1 variants show little to no increase in pelleting, indicating a failure to form higher-order species. Next, we utilized dynamic light scattering (DLS) to assess the sizes of DRP1 assemblies and aid in identifying conditions that may overcome impaired assembly. As expected by mass action, the hydrodynamic radius (R_h_) of the WT construct increased in a concentration-dependent manner before plateauing at concentrations greater than 40 µM. The G363D and G401S constructs, however, exhibited minimal size increases that plateaued at lower protein concentrations (∼5 µM, **Fig. 2B**). While both pathogenic variants demonstrated limited concentration-dependent assembly, neither construct approached the higher-order species formed by WT. These data ruled out the possibility that mutants simply possess weakened, but intact, assembly properties and demonstrate an intrinsic assembly defect.

**FIGURE 2.**
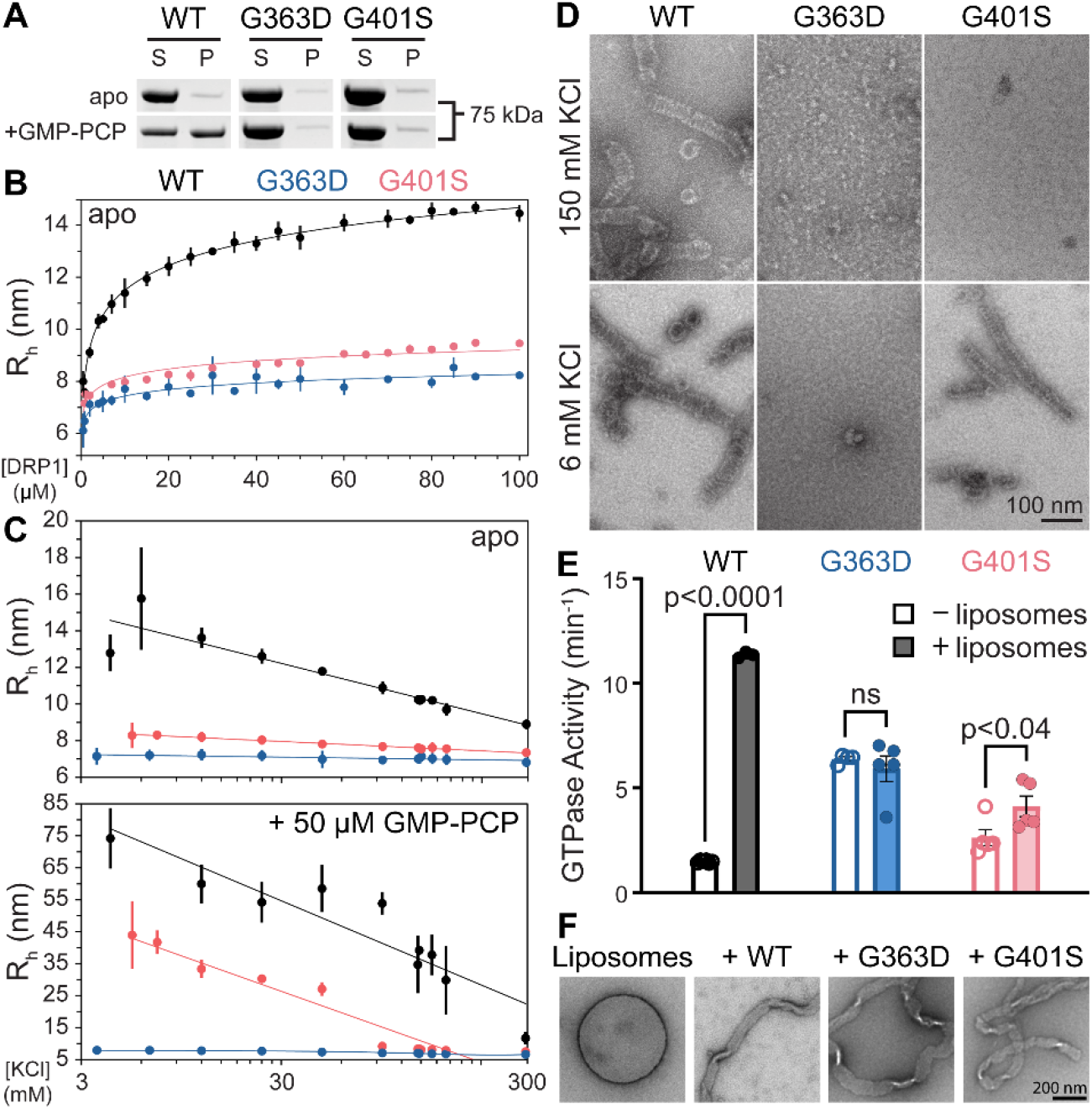
Pathogenic DRP1 variants exhibit impaired self-assembly, with G401S showing WT-like assembly only at low salt. **(A)** SDS-PAGE of supernatant (S) and pellet (P) fractions from sedimentation of 4 µM DRP1 (WT, G363D, or G401S) in the absence (top row) or presence (bottom row) of 4 mM GMP-PCP in buffer A (20 mM HEPES, pH 7.4; 150 mM KCl; 2 mM MgCl_2_; 1 mM DTT). **(B)** Hydrodynamic radius (R_h_) from dynamic light scattering (DLS) measurements as a function of protein concentration for WT (black), G363D (blue), or G401S (red) in buffer B (20 mM HEPES, pH 7.4; 150 mM KCl; 1 mM DTT) and without nucleotide. Error bars are ± SD, n = 3. **(C)** Hydrodynamic radius as a function of KCl concentration for 4 µM DRP1 WT (black), G363D (blue), or G401S (red) in the absence (top) or presence (bottom) of 50 µM GMP-PCP in buffer A (MgCl_2_ was only present alongside GMP-PCP). Error bars are ± SD, n = 3. **(D)** Representative negative-stain electron microscopy images of 4 µM DRP1 WT (left), G363D (middle), or G401S (right) with 2 mM GMP-PCP, either with 150 mM KCl (top row) or 6 mM KCl (bottom row) in buffer A. Scale bar = 100 nm. **(E)** GTPase Activity measurements for each DRP1 WT (black), G363D (blue), and G401S (red) in the absence (unfilled bars/circles) or presence (filled bars/circles) of CL-containing liposomes (40%, 35%, and 25% molar ratios of DOPC, DOPE, and CL, respectively). The reaction consisted of 500 nM DRP1, 150 μM CL-containing liposomes, 1 mM GTP, and 2 mM MgCl2 in buffer with 20 mM HEPES pH 7.4, 150 mM KCl, and 10 mM β-mercaptoethanol. All statistical tests were performed using an unpaired t-test with Welch’s correction, and p-values are indicated for each comparison (ns = not significant), n = 3 for WT and n = 5 for both G363D and G401S **(F)** Representative negative-stain electron microscopy images of 100 μM DOPS liposomes either alone (leftmost) or with one of 2 μM DRP1 WT (center-left), G363D (center-right), or G401S (rightmost) in Buffer B. Scale bar = 200 nm, n = 2 technical replicates with >50 images of each sample containing DRP1.

Prior work has firmly established that intramolecular DRP1 interactions are enhanced in lower ionic strength conditions, presumably due to increased electrostatic attractions (52, 53). Accordingly, we asked whether reducing salt concentrations would rescue the assembly of these pathogenic variants, and measured the hydrodynamic radii of DRP1 constructs across decreasing KCl concentrations (**Fig. 2C**). In the absence of nucleotide, the hydrodynamic radius of DRP1 WT displayed the expected inverse relationship between salt and size, reaching a maximum R_h_ similar to that in **Fig. 2B**. Neither G363D nor G401S showed a substantial increase in R_h_ under these conditions, indicating that these variants remain assembly-deficient without nucleotide. Upon the addition of GMP-PCP, DRP1 WT showed a nearly 5-fold increase in R_h_ that remained strongly inversely salt-dependent. G363D again showed no discernible growth in R_h_, consistent with its inability to assemble in prior assays (**Fig. 2A, B**). Strikingly, G401S exhibited a clear increase in R_h_ following the same inverse salt dependence as the WT construct, although only below physiological salt concentrations. These data indicated that G401S, but not G363D, retains assembly competence that can be revealed when both nucleotide and enhanced electrostatic interactions are present. This eliminated the possibility that both pathogenic variants can assemble normally yet require far higher protein concentrations to do so; rather, G363D appears entirely assembly-inactive, while G401S is conditionally assembly-active. To verify that the increased hydrodynamic radii reflected productive assembly instead of aggregation, we visualized these constructs using negative-stain electron microscopy (**Fig. 2D**). Under conditions with nucleotide and physiological salt, DRP1 WT readily formed rings and spirals as previously described (35– 40, 54, 55), whereas neither variant construct produced detectable assemblies. Lowering salt concentration had little effect on WT morphology and did not induce structure formation in G363D. Conversely, G401S formed WT-like rings and spirals at these low ionic strength conditions, consistent with its salt- and nucleotide-dependent increase in R_h_ (**Fig. 2C**). We next asked whether the variants could assemble upon membrane binding, which is known to stimulate DRP1 assembly and hydrolysis (17, 32, 33). As expected from prior work (32, 33), WT DRP1 increased GTP hydrolysis in the presence of cardiolipin-containing liposomes, whereas neither mutant showed a comparable membrane-dependent stimulation (**Fig. 2E**). This lack of membrane-stimulated activity was not due to a decrease in membrane binding, as both variants (and WT as expected (33, 56)) remodeled membranes (**Figs. 2F, S3**). Together, these data demonstrated that G401S, but not G363D, retains a latent capacity for self-assembly that can be revealed only under strongly permissive conditions involving both nucleotide and enhanced electrostatic interactions. Thus, neither pathogenic variant appears capable of assembly under physiological conditions.

Although G401S can assemble under strongly permissive biochemical conditions (**Fig. 2**), both pathogenic variants still form puncta in cells, suggesting that puncta do not require ordered self-assembly. To test this, we mimicked the cellular macromolecular crowding environment with PEG-8K, a common molecular crowding reagent (57–60). When recombinant DRP1 was mixed with PEG-8K, all constructs formed distinct droplets in a protein- and PEG-concentration-dependent manner (**Fig. 3**). At 1% (w/v) PEG-8K, droplet formation was minimal for all constructs. Increasing DRP1 and PEG-8K concentrations, however, produced readily detectable spherical droplets reminiscent of macromolecular and liquid-liquid phase separation (45, 46, 61–64). Increasing the concentration of DRP1 constructs to 75 µM revealed large micrometer-scale condensates and surface-wetted states (**Fig. 3A**). To investigate phase boundaries, we quantified the fluorescence intensities of labeled DRP1 constructs across gradients of DRP1 and PEG-8K concentrations (**Fig. 3B**) and examined representative fields by differential interference contrast (DIC) and fluorescence microscopies (**Fig. 3C**). DRP1 WT fluorescence increased with protein and crowder concentrations, with small condensates being abundant at 5% PEG-8K. G363D, although unable to self-assemble (**Fig. 2**), readily formed large, often fused, droplets under crowding conditions. G401S also formed condensed phases, but to an intermediate extent compared to WT and G363D. Overall, this behavior cannot be attributed to PEG-8K alone, as Ficoll PM-70 showed similar droplet formations (**Fig. S4A**). This behavior was also not a consequence of the fluorescent dye as dimeric GST did not condense under identical conditions (**Fig. S4B**). These results revealed that crowding-driven condensation is mechanistically distinct from DRP1’s canonical self-assembly properties.

**FIGURE 3.**
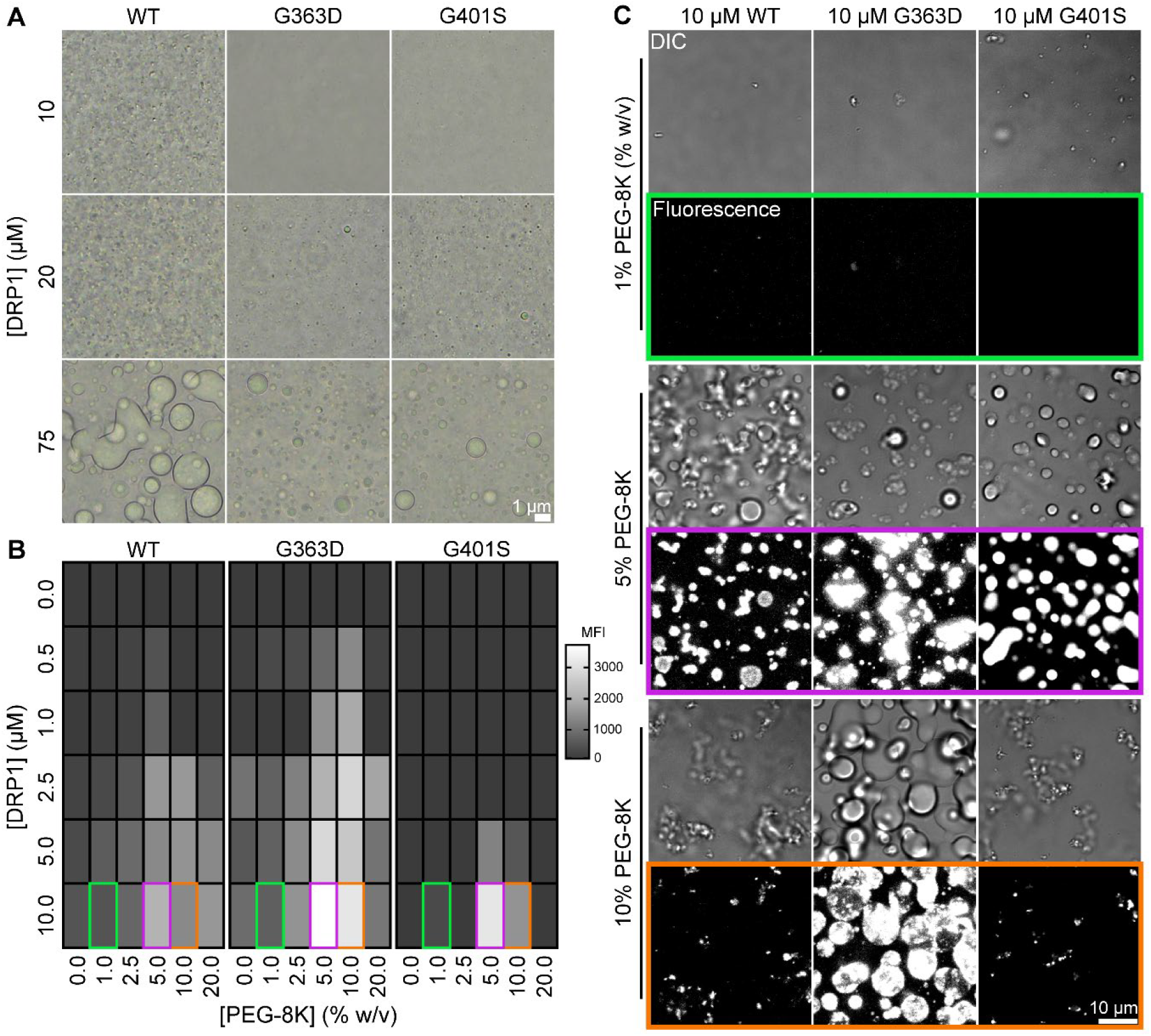
Recombinant DRP1 WT, G363D, and G401S form a continuum of condensed phases. **(A)** Representative phase contrast microscopy images of DRP1 WT (left column), G363D (middle column), or G401S (right column) at a concentration of 10 µM (top row), 20 µM (middle row), or 75 µM (bottom row) in DRP1 phase separation buffer (20 mM HEPES, pH 7.4; 150 mM KCl; 5% [w/v] PEG-8K). Scale bar = 1 µm. **(B)** Heatmap of mean fluorescence intensity (MFI) values from fluorescent images of DRP1 WT (left panel), G363D (middle panel), or G401S (right panel) at varying protein and PEG-8K concentrations in buffer (10 mM HEPES, pH 7.4; 150 mM KCl). Dark gray colored regions indicate low MFI values, whereas white regions indicate high MFI values. Regions outlined with colored boxes represent the images shown in (C). n = 9. **(C)** Differential interference contrast (DIC) and fluorescence microscopy images of DRP1 WT (left column), G363D (middle column), or G401S (right column). Displayed images depict 10 µM DRP1 with varying PEG-8K concentrations of 1% (top 2 rows), 5% (middle 2 rows), or 10% (bottom 2 rows) in buffer (10 mM HEPES, pH 7.4; 150 mM KCl). Scale bar = 10 µm, n = 9.

The concentration- and crowding-dependent properties of purified DRP1 suggest that wild-type DRP1 can access multiple condensed states distinct from well-ordered helical assembly. In prior work, we showed that the isolated Variable Domain (VD) of DRP1 is sufficient to undergo liquid–liquid phase separation under crowding conditions, implicating the VD as a potential driver of condensed-state behavior (65). These observations motivated us to ask whether DRP1 puncta in living cells exhibit dynamic and chemically sensitive properties consistent with a condensed state, and whether such behavior depends on the VD in the context of full-length protein. Expression of YFP–DRP1 produced discrete cytoplasmic puncta and larger punctate assemblies, permitting measurement of molecular exchange by fluorescence recovery after photobleaching (FRAP). Photobleaching of individual YFP–DRP1 puncta resulted in rapid fluorescence recovery, indicating continuous subunit exchange and arguing against a static, irreversibly assembled state for WT DRP1 (**Fig. 4A**). We next probed the sensitivity of these structures to chemical perturbation. Acute treatment with 1,6-hexanediol led to a marked reduction in YFP–DRP1 puncta, whereas treatment with the related control compound 1,2,3-hexanetriol had little effect, supporting selective disruption of interactions that stabilize these assemblies (**Figs. 4B, S5, S6A**). Importantly, removal of 1,6-hexanediol resulted in rapid reformation of YFP–DRP1 puncta over a period of minutes, indicating that puncta formation reflects a reversible, dynamic equilibrium rather than irreversible aggregation (**Figs. 4C, S6B**). These properties were also observed for G363D and G401S (**Fig. S6**), indicating that diol-sensitive, reversible puncta are a general property of YFP-DRP1 transiently expressed in cells, including the assembly-deficient variants.

**FIGURE 4.**
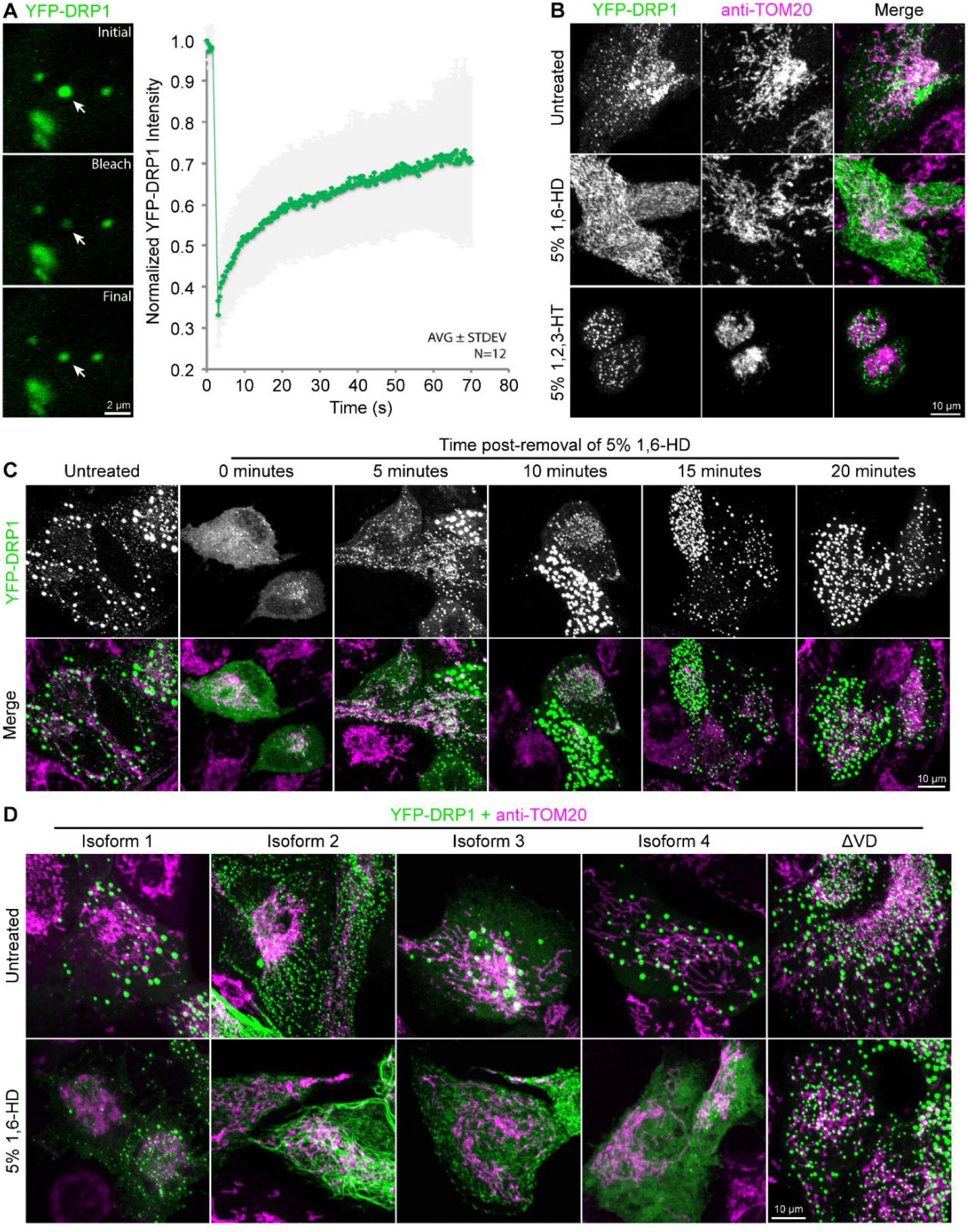
DRP1 puncta are dynamic and display reversible dissolution to 1,6-HD. **(A)** FRAP images and measurements of overexpressed YFP-DRP1 in HCT116 cells. Images were captured ever 4.28 seconds, and the resulting data were normalized, background subtracted, and bleach-corrected using pre-defined ROIs. Scale bar = 2 µm, n = 12. **(B)** Immunofluorescence images of overexpressed YFP-DRP1 in RPE cells immunostained for TOM20. Cells were treated with either 5% (v/v) 1,6-HD or 1,2,3-hexanetriol (1,2,3-HT) for 5 minutes prior to fixation. Scale bar = 10 µm, n = 3. **(C)** Immunofluorescence images of overexpressed YFP-DRP1 in RPE cells immunostained for TOM20. 5% 1,6-HD was administered to cells for 5 minutes, at which point it was removed and replaced with fresh media. Cells were fixed at the indicated times after removal of 1,6-HD. Scale bar = 10 µm, n = 3. **(D)** Immunofluorescence images of overexpressed YFP-DRP1 (of different isoforms) in RPE cells immunostained for TOM20. Cells were either untreated or treated with 5% 1,6-HD for 5 minutes prior to fixation. Scale bar = 10 µm, n = 3.

Given our prior finding that the isolated VD can phase separate *in vitro*, we anticipated that the VD would be a primary determinant of puncta formation in cells. Surprisingly, a DRP1 construct lacking the VD (ΔVD) still formed discrete puncta, despite being largely insensitive to hexanediol (**Fig. 4D**). By contrast, YFP– DRP1 isoforms that differ only in VD length and sequence due to alternative splicing formed puncta with distinct sensitivities to hexanediol. These findings indicate that puncta formation can be supported by interactions outside the VD, whereas the VD modulates the material properties and chemical sensitivity of DRP1 assemblies. This behavior is consistent with models in which intrinsically disordered regions fine-tune condensed states that are otherwise driven by additional multivalent interactions (66, 67).

Although these live-cell experiments reveal that wild-type DRP1 puncta are dynamic, reversible, and chemically sensitive, the use of overexpressed fluorescent constructs raises the possibility that puncta organization may differ from that of endogenous DRP1 (68). We therefore tested whether endogenous DRP1 puncta in control and EMPF1 patient-derived fibroblasts exhibit differential sensitivity to 1,6-hexanediol, directly assessing whether disease-associated variants alter the balance between distinct puncta states in cells (**Fig. 5A**). Untreated cells, shown for reference, recapitulated the mitochondrial morphologies described in **Fig. 1A**. Across all genotypes, DRP1 puncta are abundant at baseline, despite marked differences in mitochondrial architecture, reinforcing that puncta persistence is not tightly coupled to productive fission. Acute 1,6-hexanediol treatment then revealed striking genotype-dependent differences. In WT fibroblasts, mitochondria elongated, consistent with transient inhibition of fission (**Fig. 5B-i**), yet DRP1 puncta remained apparent and heterogeneous in brightness, with only a slight decrease in intensity and no change in total number (**Fig. 5B**). In both G363D and G401S patient fibroblasts, by contrast, hexanediol caused a pronounced, global reduction in puncta brightness: puncta remained detectable, but the bright subpopulation seen in WT cells was largely lost, resulting in a more uniform, low-intensity distribution. These differences were not explained by changes in puncta abundance or distribution, as total puncta number (**Fig. 5B-ii**) and the mitochondria-associated fraction (**Fig. 5B-iii**) were unchanged across genotypes and treatments, but instead reflected a selective change in the distribution of puncta intensities. In WT cells, puncta intensity remained broadly distributed and slightly decreased, whereas in both pathogenic variants, puncta intensities collapsed toward a uniformly dim state, as reflected by a marked reduction in intensity and intensity dispersion (**Fig. 5B-iv**). Importantly, this dispersal was reversible at endogenous levels: washout of 1,6-hexanediol restored DRP1 puncta and their intensity dispersion in both control and patient fibroblasts (**Fig. S7A, B, D**), whereas the related polyol 1,2,6-hexanetriol had little effect (**Fig. S7C, D**), supporting selective disruption of the weak interactions that stabilize these puncta.

**FIGURE 5.**
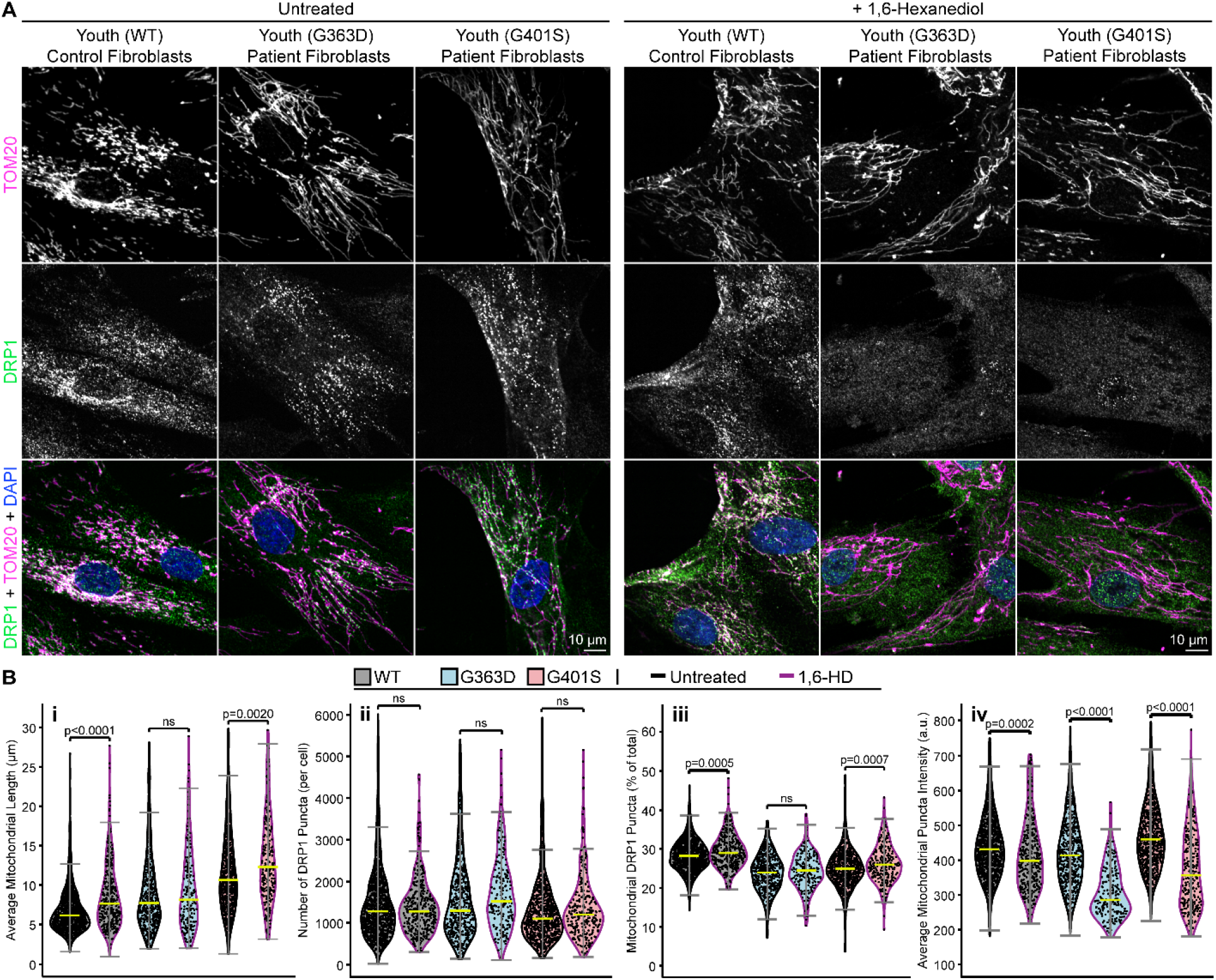
Pathogenic DRP1 variant puncta exhibit increased sensitivity to 1,6-HD compared to WT. **(A)** Representative fluorescent images of untreated human skin fibroblasts harboring wild-type DRP1 (left column) or DRP1 point mutations G363D (middle column) or G4010S (right column) that were either untreated (left panel set) or treated with 3.5% (v/v) 1,6-HD for 5 minutes (right panel set). Cells were immunostained for TOM20 (top row, magenta) and DRP1 (middle row, green), and nuclei were stained with DAPI (blue). Identical LUTs were assigned for each image. Scale bar = 10 µm. **(B)** Quantitative metrics collected from cell images of human skin fibroblasts for DRP1 WT (gray), G363D (light blue), or G401S (light red), either untreated (black outline) or treated with 1,6-HD (magenta outline). Average mitochondrial length (i) was obtained using MitoGraph software (see methods) (n = 1240, 494, or 833 untreated cells for DRP1 WT, G363D, and G401S, respectively; n = 388, 204, or 268 1,6-HD treated cells for DRP1 WT, G363D, and G401S, respectively). The total number of DRP1 puncta (ii), percentage of mitochondrial DRP1 puncta (iii), and mitochondrial DRP1 puncta intensity (iv) metrics were all obtained using the 3DSuite plugin in FIJI (see methods) (n = 1237, 494, or 831 untreated cells for DRP1 WT, G363D, and G401S, respectively; n = 388, 204, or 267 1,6-HD treated cells for DRP1 WT, G363D, and G401S, respectively). All statistical tests were performed using one-way ANOVA with Tukey’s post-hoc multiple comparisons test, and p-values are indicated for each comparison (ns = not significant). Yellow bars represent the population median. Error bars are ± 2SD. Source data for this figure are included in supplemental table **Dataset_S01**.

To further probe WT DRP1 puncta at endogenous levels, we asked whether endogenous DRP1 forms diol-sensitive puncta in other cell types. Wild-type DRP1 puncta were readily apparent in both retinal pigment epithelial and U2OS cells, and were more sensitive to diol treatment than dermal fibroblasts (**Fig. S8**). Upon removal of 1,6-hexanediol, these puncta reformed rapidly over minutes, as observed for YFP-DRP1 (**Fig. 4C, S6A**), indicating a reversible, dynamic equilibrium consistent with endogenous, wild-type DRP1 forming condensates. Collectively, these findings indicate that WT cells contain both 1,6-HD-resistant and -sensitive puncta, consistent with the coexistence of distinct puncta states with differing material properties (**Figs. S9, S10**). In contrast, pathogenic variants are biased toward the hexanediol-sensitive state, their puncta brightness and heterogeneity globally collapsed by treatment. These genotype-dependent behaviors indicate that disease-associated mutations shift the balance between puncta states in cells.

## Discussion

A central unresolved question in mitochondrial biology has been the physical nature of DRP1 puncta observed in cells. Structural and biochemical studies established that DRP1 can assemble into well-ordered oligomeric spirals and rings capable of constricting membranes, providing a compelling molecular model for mitochondrial fission (33, 34, 36, 39, 40). At the same time, live-cell imaging has consistently shown that DRP1 puncta are heterogeneous, dynamic, and frequently non-productive, with only a minority proceeding to scission during observation (20, 21, 25, 41). Puncta exchange subunits rapidly, merge and split, translocate along mitochondrial tubules, and are often observed in the cytoplasm or associated with non-mitochondrial membranes (10–12, 69). Together, these findings indicate that puncta encompass more than a single static pre-scission assembly, yet the physical basis for this heterogeneity has remained unclear.

Here, we propose a puncta continuum model in which cellular DRP1 puncta populate a spectrum of material states, ranging from more fluid, condensate-like assemblies to more ordered, fission-competent oligomers (**Fig. 6**). Endogenous DRP1 forms far more puncta than previously appreciated, with only ∼30% associated with mitochondria at steady state (**Fig. 1**). Importantly, puncta abundance and mitochondrial recruitment alone are poor predictors of fission competence: puncta persist under conditions of impaired fission, and the majority of puncta are non-mitochondrial (**Figs. 1, S1, S7A**). We probed this punctate architecture in cells using 1,6-hexanediol. While acknowledging the limitations of this reagent (70), its differential effects in our system are informative. Wild-type cells of different lineages harbor hexanediol-sensitive and -resistant puncta (**Figs. 4, 5, S6-10**), consistent with a continuum from liquid-like condensates comprised of largely unassembled DRP1 to ordered helical oligomers in a mature, scission-competent state. The documented delay between stable oligomer assembly and productive scission (21, 41) is consistent with rate-limiting maturation and conversion. Framed this way, our model extends rather than replaces the ordered-assembly paradigm: helical assemblies remain the productive fission machinery but may be reached through a condensation-associated pathway, with only a subset of puncta maturing into ordered, scission-competent assemblies. This reconciles structural models of DRP1 as a mechanochemical enzyme with the long-standing heterogeneity and non-productivity of puncta in cells. The material properties along this continuum appear tunable: DRP1 isoforms that differ in their variable domains show distinct diol sensitivities (Fig. 4D), and endogenous DRP1 puncta are more hexanediol-sensitive in RPE and U2OS cells than in fibroblasts (**Fig. S8**). We speculate that these cell-type differences may reflect variation in isoform expression, positioning the variable domain as one determinant of where DRP1 sits along the condensate–assembly continuum.

**FIGURE 6.**
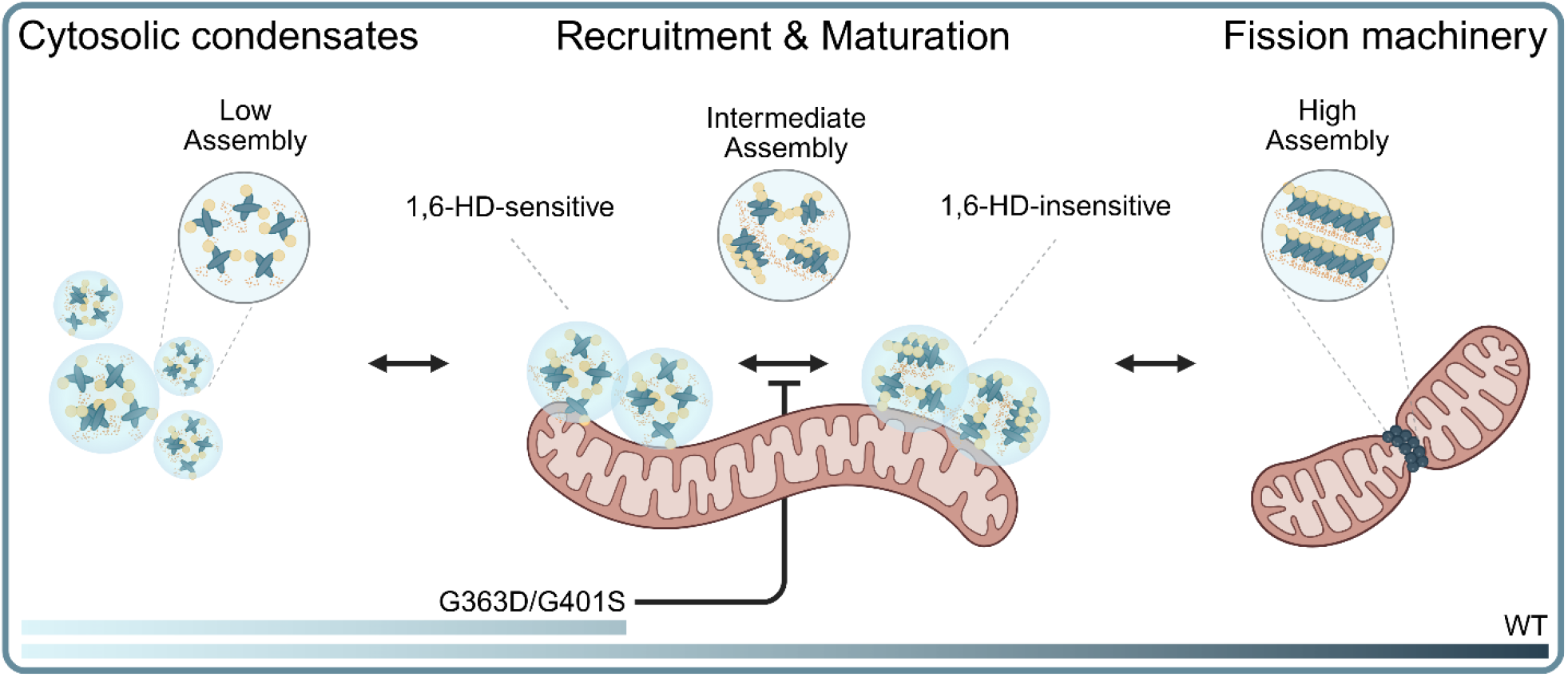
Model for DRP1 maturation from liquid-like condensates to ordered fission machinery. DRP1 forms liquid-like cytoplasmic condensates that transition through intermediate, gel-like assemblies upon recruitment to mitochondria. Wild-type DRP1 proceeds to form ordered fission rings, whereas pathogenic variants stall in less mature, 1,6-hexanediol–sensitive states, revealing a previously unrecognized maturation continuum underlying DRP1 assembly and mitochondrial fission. Created in BioRender. Hill, B. (2026) https://BioRender.com/7i6c4g8.

The heterozygous nature of EMPF1 offers a stringent test of this framework. In patient cells, wild-type and variant DRP1 are co-expressed and co-occupy shared puncta (**Figs. 1, 5, S11**). Were wild-type DRP1 to form independent, ordered assemblies at mitochondria, these would be expected to resist 1,6-hexanediol; instead, the mitochondrial-associated puncta in patient cells are uniformly diol-sensitive. The parsimonious explanation is that wild-type and variant DRP1 co-assemble, and that a single assembly-deficient subunit precludes ordered oligomerization: for a Drp1 polymer of *n* subunits that assembles at random from a 50:50 mixture of WT:variant, the fraction that is entirely wild-type decreases by ∼0.5^n^.Thus,evenfor a fission collar with tens of subunits, ordered assembly of solely wild-type species is exceedingly improbable. Wild-type DRP1 is thereby retained, together with the variant, in a shared, diol-sensitive, condensed state. These considerations indicate that wild-type DRP1 is competent to enter the condensed state in cells, while pathogenic variants are stunted and unable to further progress to ordered assembly. Because these are the mitochondria-associated puncta from which productive fission arises, this is consistent with a model in which condensation lies on the pathway to fission-competent assembly.

Pathogenic *DNM1L* variants can thus be understood as perturbations of this continuum. Despite being defective in higher-order assembly *in vitro* (15, 17), G363D and G401S form abundant puncta at endogenous levels and retain mitochondrial recruitment comparable to wild-type *DNM1L* (Fig. 1). However, upon hexanediol treatment these puncta shift globally toward a chemically sensitive, lower-intensity state and fail to maintain the brighter, hexanediol-resistant puncta observed in wild-type cells (**Fig. 5**). This shift provides a mechanistic explanation for the paradoxical coexistence of abundant puncta and profound mitochondrial elongation in patient fibroblasts: puncta formation is preserved, but progression toward a fission-competent state is impaired. The resistance of a subset of wild-type puncta is consistent with stabilization through interactions with cardiolipin (38, 71, 72) and mitochondrial adaptors (73–75), as well as with our observation that mitochondria-associated puncta are brighter even in the absence of chemical perturbation (**Fig. 1**).

Our *in vitro* data indicate that biomolecular condensation contributes to the organization of DRP1 puncta. Having previously shown that the isolated DRP1 variable domain (VD) undergoes phase separation (65), we find that full-length recombinant DRP1 accesses multiple condensed states whose properties depend on protein concentration, ionic conditions, and molecular crowding (**Fig. 3**). Both mutants condense despite remaining low-order in solution and unable to assemble (**Figs. 2, 3**), indicating that condensation does not require higher-order self-assembly; the precise oligomeric state of the species that partition into condensates, however, remains to be defined. Pathogenic variants appear to sample this condensate landscape differently: G363D remains confined to wetting condensates that fail to progress to ordered assemblies, whereas G401S is capable of condensation and limited oligomerization under permissive conditions but unable to robustly access higher-order states. These properties are consistent with structural evidence that DRP1 oligomerization requires coordinated rearrangements of conserved stalk helices and loops that mediate interface-1 and interface-3 contacts. G401 lies within the conserved L2^S^ loop implicated in interface-3, whereas G363 lies within helix α1M^S^ adjacent to the 1N^S^ loop required for these conformational transitions (39, 40). The divergent behaviors of G401S and G363D are therefore most consistent with mutation-specific perturbations of a shared stalk-mediated assembly network that bias DRP1 away from ordered polymerization and toward more condensed states (39, 40).

More broadly, our findings place DRP1 within a growing class of membrane-remodeling proteins whose organization and function are shaped by condensation. To our knowledge this is the first evidence for phase separation by a dynamin superfamily member, but related work has implicated dynamin-1 in condensate formation during synaptic vesicle recycling (76). Condensation has similarly emerged as a key organizing principle in actin assembly (77), signal tranIuction and membrane microdomain organization (78), and endocytic scission (79). These studies suggest that condensed regimes represent a general mechanism for organizing large, multivalent assemblies at membranes rather than an idiosyncratic feature of a single system.

In summary, we propose that DRP1 puncta occupy a continuum of states only a subset of which mature into fission-competent assemblies. This model reconciles disparate observations in the field and identifies biomolecular condensation as a previously unrecognized layer of DRP1 organization, regulation, and dysfunction in health and disease. Pathogenic *DNM1L* variants bias DRP1 toward immature, condensed states, decouplnig puncta formation from mitochondrial division. Our data establish a mechanistic basis for genotype-specific differences in EMPF1 and suggests that shifting DRP1 along its assembly continuum may offer a rational path toward restoring productive mitochondrial fission.

## Materials and Methods

### Cell Culture, Treatment, and Immunolabeling

Written informed consent for diagnostic molecular genetic analysis and research-based studies was obtained from all patients in accordance with the Declaration of Helsinki protocols and approved by the Newcastle and North Tyneside Local Research Ethics Committee (REC ref 2002/205). EMPF1 patient fibroblasts and associated age-matched control cells were cultured and prepared for immunofluorescence similarly as described (17). Experiments involving the expression of GFP-DRP1 were performed in HCT116 or retinal pigmented epithelial (RPE) cells as indicated, following similar culture and immunolabeling protocols as above, with an additional transient transfection step. All 1,6-hexanediol (Sigma-Aldrich; 88571-100ML-F) and 1,2,3-hexanetriol (Sigma-Aldrich; 52895-1G) solutions were combined with cell nutrient mediums to achieve their indicated concentrations and equilibrated in a sterile incubator at 37°C in an atmosphere of 5.0% CO_2_ and >90% humidity for at least 15 minutes prior to cell treatments. Cells were returned to the incubator for the indicated treatment durations. Full experimental details are provided in **Supplementary Materials and Methods**.

### Autonomous Immunofluorescence Microscopy

All microscopy images, masks, and ROIs used in analysis followed a standardized naming convention to preserve their unique identities for persistent mapping between various analysis methods and accurate merging of metrics on a per-cell basis. All cell images were acquired using a Nikon Eclipse Ti microscope base equipped with a Yokogawa CSU-W1 spinning disk confocal scanner unit (50 μm pinhole) with a Hamamatsu ORCAFlash4.0 V3 sCMOS camera. The microscope was controlled using the NIS Elements High Content package (with support for NIS.ai, GA3, and JOBS workflows), and images were captured on a 60X, 1.27 NA water immersion objective (Nikon; MRY10060). A water immersion dispenser (Nikon; MEV54006) was also used and managed within the workflows to ensure ample immersion liquid coverage of the objective throughout the imaging sessions. All images were captured with 2X averaging enabled. Fluorophores were excited sequentially with 405, 488, or 561 nanometer lasers at 50% power, and images were captured after a 200-millisecond exposure. DIC images were captured after a 1-second exposure instead. Full experimental details for image acquisition and subsequent data processing are provided in **Supplementary Materials and Methods**.

### DRP1 Expression and Purification

Recombinant DRP1 constructs were expressed in BL21(DE3) *Escherichia coli* using pET29b+ vectors encoding full-length human DRP1 (aa 1-736) fused to a C-terminal TEV cut site preceding a 6xHIS affinity tag, similarly as described (17). Full experimental details for purification and subsequent fluorophore labeling are provided in **Supplementary Materials and Methods**.

### DRP1 Sedimentation Assay

DRP1 aliquots were thawed on ice for ∼30 minutes before use. Meanwhile, β,γ-Methyleneguanosine 5′-triphosphate sodium salt (GMP-PCP) (Sigma; M3509-25MG) was freshly prepared at a final concentration of 8 mM in sedimentation assay buffer (20 mM HEPES, pH 7.4; 150 mM KCl; 2 mM MgCl_2_; 1 mM DTT). Samples were prepared by first adding protein and then GMP-PCP to sedimentation assay buffer, to achieve final concentrations of 4 μM and 4 mM, respectively. Samples were gently mixed and incubated at room temperature for 1 hour. Afterwards, 20 μL of each sample was extracted for DLS experiments. Samples were then ultracentrifuged at 26,860 xg for 45 minutes at 4°C. The supernatant was carefully removed from the pellet fraction, which was not visible by eye. The remaining pellet fraction was resuspended in sedimentation assay buffer to match the total volume of the supernatant. Supernatant and pellet fractions were added to 1X SDS-PAGE protein running buffer (25 mM Tris base, pH 8.4; 192 mM glycine; 0.1% SDS [w/v]) and incubated at 95°C for 5 minutes. Samples were then loaded in a Mini-PROTEAN TGX Stain-Free 4-20% acrylamide gradient gel (Bio-Rad; 4561096). The gel was run at 45 constant volts for ∼30 minutes or until the dye front reached the end of the stacking gel, at which point the voltage was increased to ∼120 volts and run until the dye front reached the bottom of the resolving gel. Afterwards, the gel was removed from the cassette and briefly rinsed in water before being placed in a secondary container. The gel was stained with QC Colloidal Coomassie Stain (Bio-Rad; 1610803) overnight, shaking gently at room temperature. The next day, Coomassie stain was removed, and the gel was briefly destained in water. Gels were imaged using auto-exposure timings for Coomassie-stained gels on a Bio-Rad ChemiDoc MP imaging system, with timings being slightly adjusted if needed for visual acuity.

### Dynamic Light Scattering

DRP1 aliquots were thawed on ice for ∼30 minutes before use. For protein concentration gradient experiments, DRP1 constructs were diluted to desired concentrations along the gradient in DLS assay buffer 1 (20 mM HEPES, pH 7.4; 150 mM KCl; 1 mM DTT). For salt concentration gradient experiments, DRP1 constructs were diluted to 4 μM in assay buffer 2 (20 mM HEPES, pH 7.4; 1 mM DTT) with KCl concentration being adjusted to desired concentrations along the gradient. Additionally, 50 μM GMP-PCP and 2 mM MgCl_2_ were added to DLS assay buffer 2 when assessing nucleotide-dependent assembly. Samples were briefly mixed and incubated at room temperature for 30 minutes. Meanwhile, the NanoTemper Prometheus Panta instrument was initialized, and protein concentrations (in mg/mL) and buffer components were entered into the PR.Panta Control software such that the viscosity and refractive indexes were calculated for each sample. Nearing the end of the incubation time, samples were loaded into standard capillaries (NanoTemper; PR-C002) via capillary action, ensuring no air bubbles or particulate matter were present. 2 technical replicates of each sample were included per experimental replicate. The capillary tray of the instrument was briefly cleaned with 95% ethanol and completely dried, and then samples were precisely inserted into the capillary tray. A Discovery Scan was performed on the Prometheus instrument to ensure that the samples provided ample fluorescent signal for detection and were not aggregating. The instrument was then set to perform a Size Analysis experiment at 25°C using 100% DLS laser power and 10 scans per acquisition cycle. The autocorrelation function was automatically fit in PR.Panta Analysis software from 10 sequential 5000-millisecond acquisitions to determine the hydrodynamic radius for each sample. Outlier data or failed acquisitions were removed as necessary. Data were exported to .*csv* files for further processing and plotting.

### Liposome Preparation

Lipids utilized in this study include: 1,2-dioleoyl-sn-glycero-3-phosphocholine (DOPC; Avanti; A80375), 1,2-dioleoyl-sn-glycero-3-phosphoethanolamine (DOPE; Avanti; A80725), 1,2-dioleoyl-sn-glycero-3-phospho-L-serine [sodium salt] (DOPS; Avanti; A80035), and Cardiolipin [Heart, Bovine; sodium salt] (CL; Avanti; A84012). All lipids were suspended in chloroform and dried under nitrogen gas and SpeedVac for 30 minutes at room temperature. Afterwards, lipids were resuspended in liposome buffer (20 mM HEPES, pH 7.4; 150 mM KCl; 1 mM DTT) to reach a stock concentration of 2 mM. CL-containing liposomes (40%, 35%, and 25% molar ratios of DOPC, DOPE, and CL, respectively) were extruded using the Mini Extruder Pro Kit with base (Avanti; A81411) through 0.4 μM PC Membranes (Avanti; A89040) at 37°C at least 11 times. DOPS liposomes were freeze-thawed twice in liquid nitrogen instead of being extruded.

### Negative Stain Electron Microscopy

Recombinant DRP1 constructs were thawed on ice for ∼30 minutes before use. For nucleotide experiments, proteain was diluted to 4 μM in assay buffer (20 mM HEPES, pH 7.4; 2 mM GMP-PCP; 2 mM MgCl_2_; 1 mM DTT). KCl concentrations were either 6 or 150 mM, depending on the experiment. For liposome experiments, protein was diluted to 2 or 4 μM in liposome buffer (20 mM HEPES, pH 7.4; 150 mM KCl, 1 mM DTT) containing either 100 μM DOPS or 150 μM CL-containing liposomes (see **Liposome Preparation**). Solutions were allowed to equilibrate at room temperature for 30 minutes prior to beginning the experiment. 5 μL of each sample was added to a pre-discharged Formvar/Carbon 300-mesh grid (Electron Microscopy Sciences; FCF300-CU) and incubated for 5 minutes. Liquid was removed with Whatman Grade 1 filter paper from the edges of the grid. The grid was rinsed with 10 μL ddH_2_O for a maximum of 20 seconds, with liquid removed as above. 10 μL of filtered 2% uranyl acetate solution (Electron Microscopy Sciences, 22400) was added to the grid for a maximum of 20 seconds, with liquid removed as above. Grids were inserted into a JEOL JEM-120i, 120kV TEM instrument and imaged at 10,000X, 25,000X, and 80,000X magnifications.

### Phase Separation Assays

Individual aliquots of each of the DRP1 constructs (WT, G401S, G363D), both labeled and unlabeled, were thawed on ice prior to use in phase separation assays. Labeled aliquots were protected from light during thawing. A #1.5 high-performance cover glass 96-well plate (Cellvis; P96-1.5H-N) or a Costar 96-well clear flat bottom plate (Corning; 3909) was used for each experiment. HEPES buffer pH 7.4, KCl, and ddH_2_O were added to all wells at appropriate volumes to reach 10 mM HEPES and 150 mM KCl, targeting a final volume of 100 μL. The volumes of these reagents were calculated to offset contributions of HEPES and KCl present in the labeled and unlabeled protein stocks. PEG-8K (diluted in ddH_2_O) was then added to the appropriate wells to reach the desired percentage (w/v). Lastly, labeled and unlabeled protein was added to each well prior to careful yet ample mixing to avoid introducing air bubbles that could obfuscate light scattering measurements. In fluorescently labeled experiments, Cy5-DRP1 WT and Cy5-DRP1 G363D accounted for 20% of the total protein concentration in the wells, and AlexaFluor568-DRP1 G401S accounted for 5% of the total protein concentration in the wells. The plate was then incubated at room temperature and protected from light prior to downstream data collection.

### Fluorescence Microscopy

Fluorescence microscopy and differential interference contrast microscopy were performed on a Nikon A1R laser scanning confocal microscope equipped with an A1-DUG Hybrid GaAsP/PMT detector. The microscope was controlled using NIS-Elements Confocal & Enhanced Resolution software package (versions 5.0 and newer), and images were captured on a 60X, 1.40 NA oil objective. 3 random locations within each condition were imaged. For DRP1 WT and G363D constructs labeled with Cy5-Azide, fluorescence was stimulated with a 640 nm laser at 15% power, -127 offset, and 125 gain. Images were captured after a 1-second dwell time. DIC images were captured using the transmitted detector via the 647 nm laser, with 0 offset and 80 gain. For DRP1 G401S labeled with AlexaFluor 568, fluorescence was stimulated with a 561 nm laser at 10% power, -127 offset, and 50 gain. Images were captured after a 1-second dwell time. DIC images were captured using the transmitted detector via the 561 nm laser, with 0 offset and 80 gain. All images were collected at room temperature at least 1 hour after experiment setup.

### Fluorescence Microscopy Image Analysis

Analysis of fluorescence microscopy images was performed with ImageJ (FIJI; versions 1.52n and newer). A custom script was written to batch-process the images and output results to .*csv* files. Images were split into their individual component channels, with the fluorescent channel being used to generate a maximum intensity projection, whereas the DIC channel was not processed further. The maximum intensity projections were used to collect various metrics, such as image area, mean fluorescence intensity, standard deviation of mean fluorescence intensity, and integrated density across the entire image field. The images were then thresholded both above and below a pixel intensity value of 400, such that fluorescence within objects and/or the background could be independently evaluated with the same list of metrics mentioned above. Partition coefficients were calculated using the integrated density within the thresholded objects divided by the total integrated density of the entire image field. Data points were then plotted as heatmaps as a function of DRP1 and PEG-8K concentrations in R.

### YFP-DRP1 FRAP Microscopy

Full experimental details for cell culture and transfection are provided in **Supplementary Materials and Methods**. The microscope environmental chamber was pre-equilibrated to 37°C, 5% CO2, and ∼90% humidity prior to staging of the dish. FRAP was achieved using a Zeiss LSM 510 laser point-scanning confocal microscope and 100X 1.45 NA objective. A 511x200 pixel region was imaged on the objective (3.5 zoom). A 50x50 pixel square ROI was used to define the bleach region, and was photobleached over 20 iterations using a 488 nm laser at 100% laser power. 8 images were captured prior to the photobleaching event, and 292 images were taken after photobleaching. Images were captured once every ∼4.28 seconds. Additional 50x50 pixel square ROIs were defined for background fluorescence and non-specific photobleaching, following the same criteria above. The resulting data were normalized, background subtracted, and bleach-corrected using each ROI.

### YFP-DRP1 Immunofluorescence Microscopy

Full experimental details for cell culture, transfection, and immunolabeling are provided in **Supplementary Materials and Methods**. Images were acquired using a Nikon Eclipse Ti microscope base equipped with a Yokogawa CSU-W1 spinning disk confocal scanner unit (50 μm pinhole) with a Hamamatsu ORCAFlash4.0 V3 sCMOS camera. The microscope was controlled using the NIS Elements High Content package (with support for NIS.ai, GA3, and JOBS workflows), and images were captured on a 60X, 1.42 NA oil immersion objective (Nikon; MRD71670). All images were captured with 2X averaging enabled. Fluorophores were excited sequentially with 488 and 561 nanometer lasers at 50% power, and images were captured after 1-second or 500-millisecond exposures, respectively.

## Custom Code

All custom code for data acquisition, analysis, and visualization is available for download at https://github.com/Hill-Lab/DRP1_Puncta_Analysis or 10.5281/zenodo.21969344.

## Acknowledgements

We would like to acknowledge the Children’s Research Institute and Cardiovascular Research Center at the Medical College of Wisconsin (MCW) for allowing us to use their microscopy systems to support this work. We thank the Research Computing Centers at MCW and CU Boulder for access and use of their high-performance computing clusters for large-scale data processing and analyses. This study was also supported by the Electron Microscopy Facility at CU Anschutz (RRID: SCR_025521), and data were collected using the JEOL JEM-120i, 120 kV TEM supported by NIH grant 1S10OD036258-01.

Funding for this study was provided by the following sources: National Institutes of Health grants R01GM067180 and S10 OD025036 (RBH). RWT was supported by the Wellcome Centre for Mitochondrial Research (203105/Z/16/Z), the Mitochondrial Disease Patient Cohort (UK) (G0800674), the Medical Research Council (MR/W019027/1), the Lily Foundation, the UK NIHR Biomedical Research Centre for Ageing and Age-related disease award to the Newcastle upon Tyne Foundation Hospitals NHS Trust, LifeArc and the UK NHS Highly Specialised Service for Rare Mitochondrial Disorders of Adults and Children. MO was supported by the Fight for Sight and the Academy of Medical Sciences. EHRM was supported by a National Institute of General Medical Sciences (NIGMS) Postbaccalaureate Research Education Program (PREP) R25 Grant #R25GM140243 with additional support from the University of Colorado (CU) Comprehensive Cancer Center (GRANT# P30CA046934), CU School of Medicine, Department of Surgery and Division of Medical Oncology, and the Skaggs School of Pharmacy and Pharmaceutical Sciences. MJF was supported by National Institutes of Health grant T32GM149361. The content is solely the responsibility of the authors and does not necessarily represent the official views of any of the aforementioned funding agencies.

Portions of this text were improved by ChatGPT v5.2.

