## Supplementary material for "Pathogenic DRP1 variants reveal a role for biomolecular condensation in mitochondrial fission": supp_info

**This PDF file includes:**

Supplementary Materials and Methods

Figures S1 to S11

SI References

### Supplementary Materials and Methods

#### EMPF1 Patient Fibroblast Culture, Treatment, and Immunolabeling

EMPF1 patient fibroblasts and associated age-matched control cells were grown in Dulbecco's Modified Eagle Medium:Nutrient Mixture F-12, HEPES (Gibco; 11330-032) supplemented with 10% (v/v) FBS, 1X non-essential amino acids (Gibco; 11140-050), 50 μg/mL uridine, 50 U/mL penicillin, and 50 μg/mL streptomycin (Gibco; 15140-122). Cells were incubated at 37°C in an atmosphere of 5.0% $CO_2$ and >90% humidity. Cells were plated onto #1.5 high-performance cover glass 96-well (Cellvis; P96-1.5H-N) or 24-well (Cellvis; P24-1.5H-N) plates and grown for 48 hours prior to treatment at the incubation conditions described above. Approximately 10,000 and 15,000 cells were initially seeded in each well of the 96- and 24-well plates, respectively. 1,6-hexanediol (Sigma-Aldrich; 88571-100ML-F) was diluted in media to achieve the desired 3.5% (v/v) solution. These solutions were then allowed to equilibrate within the incubator for at least 15 minutes before treatment. Media was aspirated from all wells, proceeding one column at a time. Untreated wells received fresh media, and treated wells received the appropriate 1,6-hexanediol:media solution for the allotted incubation time. The cells were returned to the incubator for the duration of treatment. The cells were protected from light throughout all incubation steps after this point. Following treatment, media and treatment solutions were aspirated from all wells, proceeding one column at a time. Cells were fixed in 4% paraformaldehyde (Thermo; 043368-9M) in 1X PBS (Gibco; 70011-044) for 25 minutes, shaking gently at room temperature. Next, cells were permeabilized in 0.15% (v/v) Triton X-100 (Sigma-Aldrich; X100-1L) in 1X PBS for 15 minutes, shaking gently at room temperature, then blocked in 3% (w/v) BSA (Sigma-Aldrich; A7030-100G) and 0.3% (v/v) Triton X-100 in 1X PBS for 1 hour, shaking gently at room temperature. Primary antibodies, mouse anti-DRP1 (BD Biosciences; 611113) and rabbit anti-TOM20 (Santa Cruz; sc-11415), were diluted 1:100 in blocking buffer and added to the cells overnight, with gentle rocking at 4°C. Afterward, cells were washed with 1X PBS 3 times for 5 minutes each before receiving the appropriate conjugated secondary antibody, either goat anti-mouse AlexaFluor 488 (Life Technologies; A-11001) or goat anti-rabbit AlexaFluor 568 (Life Technologies; A-11011), diluted 1:500 in blocking buffer for 1 hour, rocking gently at room temperature. Subsequently, cells were washed again with 1X PBS 3 times for 5 minutes each, then received DAPI solution (Thermo; 62248) diluted 1:1000 in blocking buffer for 10 minutes, rocking gently at room temperature. Cells were washed in 1X PBS 3 times for 5 minutes each, then stored in 1X PBS containing 0.2% (w/v) sodium azide (Thermo; S2002-500G), sealed tightly in parafilm, and stored at 4°C until they were imaged.

#### Autonomous Immunofluorescence Microscopy (continued from Main Text)

Custom Nikon JOBS and GA3 workflows were created to autonomously collect images, which are available at 10.5281/zenodo.21969344 (although these will only work on Nikon systems that have the required software upgrades to run them). Autonomous image acquisition began with defining the precise well-plate dimensions as per the manufacturer's specifications (many plate varieties are also predefined within the software). Once entered, the custom plate was selected in the software. Next, it was set on the microscope stage to have its position calibrated in the software for precise autonomous movements. The wells to be imaged were then selected, at which point the optimal movement path was calculated (in this case, snaking across all wells, column by column). The water immersion dispenser was primed for use and set to dispense liquid on the objective lens. The stage was manually moved such that the first well to be imaged in the path was centered above the objective lens, and focus was manually adjusted until the base focal plane was set (the software needs help in setting initial focus for later steps to proceed smoothly).

The next series of instructions was followed for each well until valid end criteria were met (described later). First, the acceptable imaging boundaries of each well were restricted to a toroid, such that positions too near the edges or center were invalid (this was highly customizable). Within the acceptable toroidal region, a set of 40 positions was randomly generated as potential imaging destinations (pre-defined positions and various randomization algorithms can also be utilized here). The system would then proceed to each defined point in the order of its generation (instead of optimal path). Once the microscope stage moved to an imaging point, the Nikon AutoFocus and PerfectFocus systems were initiated to define a new positional focus. In any instance where internal system checks used in defining focus failed, the focus was returned to the user-defined position, and focusing strategies were re-performed. If this still failed to pass internal system checks, the point was skipped. If successful, the microscope would take a quick snapshot of the region stimulated with the 405 nm laser. This image would be the input to a GA3 helper script, in which nuclear DAPI signal above a specified threshold would then be counted as a “cell” for detection purposes. If no nuclei were detected, the point was skipped. If many nuclei (>8) were detected, the microscope would continue to a later step in the procedure for final image acquisition. Otherwise, if nuclei detection was successful but not considered as many (>1 and <8), the stage would move to the center of each detected nucleus and obtain another quick snapshot. The same nuclei detection script was rerun on this new image, and the number of detected nuclei was recounted and compared to the original. If the new image had greater counts of nuclei than the previous, the stage position of the original random point was offset to the position of the new image containing more “cells”. In addition to raw counts, the area fraction of the image occupied by nuclear signal was also analyzed. In the event that several nuclei positions yielded the same total count of nuclei, the snapshot that had the greatest area fraction of these nuclei in frame was favored for final acquisition. This additional metric aided in the reduction of final acquisitions having many “cells” on the edge of the image series when another offset position may have the same raw counts with more “cells” visibly in frame. This optimization sub-protocol was performed until each original central nucleus position was tested, with the result being the image containing the greatest number of “cells” that best filled the capture region near the original randomly generated point. Following the initial detection of many “cells” and/or optimization of the image field to increase “cell” count and area fraction, the system began its final image acquisition sequence. The refocused plane was set as the home focus position during acquisition, at which 1 image was taken, and relative to which 10 images were taken below and 17 images were taken above, with a Z-step height of 0.3 µm between each slice (28 slices total). At each slice, an image was collected for each desired wavelength, which in this instance were 405, 488, and 561 nm (for DAPI, anti-DRP1, and anti-TOM20 signal, respectively). See the first paragraph in this section for precise imaging parameters for each channel. Once the acquisition was complete, the image was saved with information detailing the well position of the plate and the specific point number in sequence. The protocol would then loop to the next point in sequence within a well and repeat all outlined steps. The water immersion dispenser would remove excess water and reapply liquid every 2 wells (this was highly customizable).

For the system to proceed to the next well to be imaged, several criteria needed to be met, all of which were defined prior to running the automation scripts. Specifically, the system would maintain the combined sum of the number of nuclei detected for each valid point where final image acquisition was initiated. Additionally, a minimum number of final acquisitions was set for each well. As such, at least 25 total “cells” were imaged in at least 7 image series per well, per plate.

**Immunofluorescence Microscopy Single Cell Segmentation**

Following the autonomous acquisition, images were subjected to another custom Nikon GA3 script for the autonomous production of single-cell masks, which are available at [10.5281/zenodo.21969344](10.5281/zenodo.21969344). First, an image was split into its individual component channels, of which only the 405-nanometer laser and DIC channels were used. The nuclear DAPI fluorescence that resulted from excitation via a 405-nanometer laser was first subjected to a Gaussian blur filter, such that the entire organelle could be easily identified as a single object via homogeneous area detection. If multiple nuclei were found to be directly adjacent to one another, an additional algorithm was employed to define them as separate objects (based on differences in signal intensity). By this point, all nuclear DAPI signals had been identified as unique objects and were preserved as such. Meanwhile, the DIC channel had its contrast distorted and enhanced through several algorithms in sequential steps, such that the dark cell boundaries became very accentuated and easily discernible. Homogeneous area detection was then implemented to identify the regions surrounded by the boundary signal as single objects, similar to that performed on the DAPI signal. Next, object math was performed between the unique nuclear objects and cell boundary objects, such that only the cell boundaries that contained at least one nuclear object were preserved. While the combination of prior steps was successful in identifying cellular regions, it often struggled to disambiguate cells in extremely close proximity where cell borders were slightly ambiguous. As such, additional steps were implemented to handle these scenarios. First, the unique nuclear objects underwent a watershed algorithm while preserving the context of the cellular boundaries. These nuclear watershed objects were combined with the original cell boundary objects, which resulted in the distinct separation between nearby cells. Lastly, these new cell objects were uniquely identified and converted into a binary mask.

The resulting masks generated in the above process were sequentially opened in ImageJ alongside the original images from which they were generated. The masks were thresholded above zero intensity to generate an ROI for each cell within the image, which were sequentially indexed and saved following the same naming conventions as other image files to preserve their unique identity. ROIs were then individually looped through for each image, from which all data and metadata for each channel and z-slice were copied (aka cropped) into a new image file and saved independently following the same conventions. From this, each cell could be individually evaluated with various analyses to obtain single-cell metrics for each cell within the dataset.

**Immunofluorescence Microscopy Batch Image Analysis**

Several analysis workflows were initiated immediately following the generation of cropped single-cell images as outlined above. All scripts used in this pipeline are available at 10.5281/zenodo.21969344. First, each cropped cell image was split into its constituent channels, of which only the anti-DRP1 (488 nm) and anti-TOM20 (561 nm) signals were used.

For collecting MFI and sdMFI data, the native FIJI function “Set Measurements…” was called with the following additional arguments: "area mean standard min perimeter integrated median area_fraction stack limit display redirect=None decimal=4". Next, the DRP1 signal for a given cell was thresholded above 150 (determined to be just above the background signal noise floor), and the native FIJI function “Measure” was called. The resulting data were saved as *.csv* files and imported into R for merging between replicates, generation of plots, and statistical analyses.

For obtaining Pearson’s and Mander’s colocalization coefficients between the DRP1 and TOM20 signal, the “coloc2” function in FIJI was called. In this function call, the DRP1 signal was always specified as channel 1, and the TOM20 signal was always specified as channel 2. Moreover, the analysis was limited to the ROI of the cropped cell. Additional arguments were added to specify the use of a bisection threshold regression strategy, using a point spread function with a value of 3, and implementing Costes’ randomizations with a value of 10. The resulting data were saved as *.csv* files and imported into R for merging between replicates, generation of plots, and statistical analyses.

For generating masks, the TOM20 signal was thresholded in three dimensions above and below an intensity value of 600 for each cell. This value was determined to be above the noise floor and background TOM20 signal (if at all present) while also preserving the majority of the positive TOM20 signal. Mitochondrial masks were specifically generated from the signal above the threshold (positive mask), while the cytoplasmic masks were generated from the signal below the threshold (negative mask). After their generation, the raw DRP1 signal was duplicated twice, with each duplicate being divided by either the positive or negative TOM20 mask via the “imageCalculator” function in FIJI. This produced new images containing all the DRP1 signal within their respective masked region while omitting all DRP1 signal outside that same region. These outputs were then multiplied by their same masks to return the raw pixel intensities to their original values (as they were reduced during the division step). The resulting images were then used in mitochondrial or cytoplasmic DRP1 puncta analysis.

For individual DRP1 puncta analysis, the 3DSuite plugin was installed, which enabled three-dimensional segmentation and conservation of particles between slices (1). First, the "3D Iterative Thresholding 2 (beta)" function was called on each masked image from above with the following parameters: a minimum pixel volume of 8, a maximum pixel volume of 500, a minimum signal threshold of 150, a minimum contrast of 0, the criteria method MSER, the threshold method STEP, segment results being All, and a value method of 10. The beta version of the function was used as it supported the greater bit-depth images produced by the image arithmetic in the masking steps. The pixel volume limits were set to prevent the inclusion of potential single-particle noise on the small end and aggregate puncta on the large end. The intensity threshold was determined to be just above the noise floor of the DRP1 signal. Many other permutations of criteria and threshold methods were possible via this function, all of which were first tested on a small subset of the data, and from which it was determined that the combination of MSER and STEP produced the best segmentation results. All other parameters that were not discussed had their default values used. After completion of the function call, a new image stack was created containing the objects detected during the segmentation and thresholding, condensed into a single plane. Each image stack had 3 of these planes, with the detected object size increasing in each. The last plane rarely contained any objects, so it was omitted. The first two planes were subjected to XOR image arithmetic, such that objects were preserved in the result if and only if they existed in exactly one of the two planes. This was done to prevent potential aggregate objects in the second plane from overlapping with multiple smaller objects in the first plane. It's worth mentioning that the resulting dataset was highly similar when using AND or OR image arithmetic instead of XOR, but the latter was chosen since it appeared to slightly reduce the presence of overly large objects. The result of the XOR image arithmetic was then used as an input to the "3D Intensity Measure" function in 3DSuite, from which intensity data was generated for every individual punctum object identified in the cell, for both sets of masks. The resulting data were saved as *.csv* files and imported into R for merging between replicates, generation of plots, and statistical analyses.

**MitoGraph Batch Analysis**

MItoGraph was used to quantify mitochondrial morphology as described (2). In brief, mitochondria were first segmented from the TOM20 signal of autonomously cropped single cells as outlined prior. The resulting cells were individually visually inspected for accuracy and were grouped into folders containing 50 images each (for parallel processing). These folders were then uploaded to Linux-based HPC servers that housed the MitoGraph software. MitoGraph was executed from its local container within a terminal via the following: "run_mitograph -xy 0.11 -z 0.3 -adaptive 10". This specified the image scale as well as the thresholding strategy for parsing the mitochondrial signal. An additional "-path" argument was supplied to specify the location of the uploaded data folders. Following the completion of MitoGraph processing, the results were further inspected for accuracy, with any outliers being automatically removed via subsequent filtering and denoising following a Gaussian fit model. Several R scripts were used to parse the resulting metrics, such as the MitoGraph Connectivity Score and Average Degree of Branching. "The Average Mitochondrial Length" metric (as reported in figures) was derived from dividing the "Total Mitochondrial Length" parameter by the "Total Connected Components" parameter, both of which were directly computed from MitoGraph. This more accurately reflects the average length of combined mitochondria as they appear in mitochondrial networks, rather than that of a single mitochondrion (the length of which does not drastically differ as reported by MitoGraph). Unique individual cell identifiers were carefully preserved to ensure accurate merging of quantitative metrics obtained between replicates and via other analyses and are uploaded in supplemental table **Dataset_S01**. Data were plotted with R, and statistical analyses were performed using one-way ANOVA with Tukey's post-hoc comparisons. Data processing and plotting scripts are available at 10.5281/zenodo.21969344.

**DRP1 Expression and Purification** (continued from Main Text)

Recombinant DRP1 constructs were expressed in BL21(DE3) *Escherichia coli* using pET29b+ vectors encoding full-length human DRP1 (aa 1-736) fused to a C-terminal TEV cut site preceding a 6xHIS affinity tag. Transformed cells (from glycerol stocks) were grown in LB broth (Miller) cultures containing 30 mg/mL kanamycin, shaking at 220 rpm at 37°C overnight (~16-18 hours). Starter cultures were used to inoculate larger 1 L growths in super broth (RPI; S33055-5000.0) also containing 30 mg/mL kanamycin, shaking at 220 rpm at 37°C until $OD_{600}$ measurements were between 1.2 and 1.8 (usually between 2.5 and 4 hours). Once within the desired $OD_{600}$ range, cultures were induced with 1 mM isopropyl 1-thio-β-D-galactopyranoside (IPTG) (RPI; I56000-25.0) and were returned to the incubator to shake at 180 rpm at 18°C overnight (~16-18 hours). Afterwards, cultures were collected and centrifuged at 3,789 xg for 30 minutes at 4°C to harvest cell pellets, which were resuspended in Buffer A (20 mM HEPES, pH 7.4; 500 mM KCl; 40 mM Imidazole; 0.02% [w/v] sodium azide) and frozen at -80°C until proceeding with purification.

Cell pellets were fully thawed in room-temperature water, at which point a half tablet of Complete™, EDTA-free Protease Inhibitor Cocktail (Sigma; 11873580001), 100 µg/mL lysozyme (OmniPur; 5960), ~1µg of DNAse I (Sigma; 10104159001), and 12 µM of $CaCl_2$ and $MgCl_2$ were added to the pellets. These were then gently mixed and incubated on ice for up to 30 minutes. Pellets were sonicated on ice at ~50% power continuously for 15 seconds, and were then rested for 45 seconds. These steps were repeated until a total of 4 minutes of sonication time had elapsed. Pellets were then centrifuged at 184,000 xg for 1 hour at 4°C to obtain the clarified lysate.

The clarified lysate was passed through 0.45 µm PVDF syringe filters (Millipore) and loaded onto a His60 Nickel Superflow Resin (TaKaRa Bio; 635662) column that had been pre-equilibrated with Buffer A using an AKTA FPLC. The column was washed with 10 column volumes of Buffer A, then with 10 column volumes of Buffer A containing 1 mM ATP, then with 10 column volumes of Buffer A containing 0.5% (w/v) CHAPS (Sigma; C3023-100G), and then with an additional 10 column volumes of Buffer A. Next, the protein was eluted from the column along a gradient of Buffer B (20 mM HEPES, pH 7.4; 500 mM KCl; 500 mM Imidazole; 0.02% sodium azide) and collected in 3 mL fractions.

Fractions containing protein (based on UV chromatograms and qualitative Bradford assay) were combined (usually between 15-20 fractions), to which 1% (v/v) HyperTEV60 (3) was added. This mixture was then dialyzed into DRP1 dialysis buffer 1 (20 mM HEPES, pH 7.4; 150 mM KCl; 0,1% [v/v] BME) for ~4 hours at 4°C, stirring gently. Nearing the end of dialysis, a diethylaminoethyl (DEAE) Sepharose Fast Flow (Cytiva; 17070901) column was prepared and charged with 10 column volumes of 2 M NaCl. The DEAE column was attached below a nickel affinity column in series, at which point both columns were equilibrated with 10 column volumes of reverse-nickel buffer (20 mM HEPES, pH 7.4; 150 mM KCl; 0.02% [w/v] sodium azide) using an AKTA FPLC. The eluted fractions were removed from dialysis and loaded onto the tandem columns. Cleaved protein was collected in the flow-through upon washing with 10 column volumes of reverse-nickel buffer, which was then dialyzed into DRP1 dialysis buffer 2 (20 mM HEPES, pH 7.4; 150 mM KCl; 1 mM EGTA; 0.1% [v/v] BME) for ~4 hours at 4°C, stirring gently. Afterwards, the protein was dialyzed into DRP1 dialysis buffer 3 (20 mM HEPES, pH 7.4; 150 mM KCl; 1 mM DTT) overnight (~16-18 hours) at 4°C, stirring gently.

The protein was then removed from dialysis and passed through 0.45 µm PVDF syringe filters. Next, the protein was loaded into a 30 kDa MW cut-off concentration column (Millipore) and was centrifuged at 3,300 xg at 4°C in 30-minute intervals. Between runs, the flow-through was discarded, and additional protein was added to the column as necessary. This continued until the entirety of the protein sample was concentrated to a volume of ~1-5 mL. Protein concentration and purity were determined using the theoretical extinction coefficient and the absorbance at 280 nm using measurements obtained from a NanoDrop OneC Microvolume UV-Vis Spectrophotometer (Thermo). The purified protein was then distributed into single-use aliquots (volume varied), flash frozen in liquid nitrogen, and stored at -80°C until use.

**DRP1 GTP Hydrolysis Assay**

DRP1 GTPase activity was measured using malachite green to detect free phosphate as described previously (4). Briefly, DRP1 proteins, with or without liposomes, were incubated for 15 minutes at room temperature without GTP or MgCl2, and were moved to 37°C for 5 minutes prior to initiating the reaction via addition of GTP and MgCl2. The reaction consisted of 500 nM DRP1, 150 μM 25% CL-containing liposomes, 1 mM GTP, and 2 mM $MgCl_2$ in buffer with 20 mM HEPES pH 7.4, 150 mM KCl, and 10 mM β-mercaptoethanol. Aliquots of the reaction were taken every 5 minutes for 30 minutes and added to EDTA (100 mM final concentration) on ice. The amount of free phosphate at each time point was then measured by adding a malachite green solution (1 mM malachite green, 10 mM ammonium molybdate, 1 N NaOH) and measuring absorbance at 640 nm after 10 minutes (Cytation5, BioTek). The initial rate of free phosphate generated was then used to calculate a $k_{obs}$ based on the DRP1 concentration tested.

**DRP1 Cy5 Labeling**

DRP1 WT and G363D were labeled with a Cy5-azide fluorophore on methionine residues via copper click chemistry on a sulfamide conjugate intermediate. To begin, a 10-fold molar excess of propargyl oxaziridine was added to each freshly prepared protein construct in buffer (20 mM HEPES, pH 7.4; 150 mM KCl; 2 mM $MgCl_2$; 1 mM BME; 0.02% [w/v] NaAz) and incubated at room temperature for 15 minutes. The reaction mixtures were then added to Sephadex G-25 M PD-10 desalting columns (GE) that were pre-equilibrated with the same buffer as above, from which 1 mL fractions were collected, eluting with 1 mL of buffer each time. These fractions were combined and concentrated to a volume of ~300 μL in a 30 kDa MW cut-off concentration column (Millipore), centrifuging at 3300 xg and 4°C. Final concentrations of 25 mM HEPES and 40 μM Cy5-azide dye (Click Chemistry Tools, now VectorLabs) were achieved after their addition to the entirety of each of the concentrated protein samples. 1.25 mM BTTAA and 250 μM $CuSO_4$ were combined and vortexed until light blue in color, at which point 12.5 mM sodium ascorbate was added and further vortexed until clear. This solution was added to each of the concentrated protein samples and incubated at room temperature for 10 minutes, protected from light. Afterwards, the samples were applied to a second set of pre-equilibrated PD-10 columns, collected in 1 mL fractions as above, and combined and concentrated to a volume of ~1 mL in a 30 kDa MW cut-off concentration column. Protein concentration and degree of labeling were determined with equations 1 and 2 using measurements obtained from a NanoDrop $One^C$ Microvolume UV-Vis Spectrophotometer (Thermo). DRP1 WT and G363D (15 methionines) had degrees of labeling of ~0.44 and ~0.51 dye molecules per protein molecule, respectively. Labeled proteins were aliquoted into single-use 20 μL samples and frozen at -80°C until use.

**DRP1 AlexaFluor 568 $C_5$ Maleimide Labeling**

DRP1 G401S aliquots frozen at -80°C were thawed on ice and dialyzed overnight (~16 hours with a buffer swap at ~10 hours) into 20mM HEPES pH 7.4, 150mM KCl, 2mM $MgCl_2$, and 1 mM TCEP at 4°C. This step was needed to remove the DTT from the buffer system leftover from purification that would otherwise interfere with maleimide conjugation of the fluorophore. Protein was then removed from dialysis and filtered through a 0.45 μm PVDF syringe filter (Millipore) to remove potential aggregates and other large particles introduced via dialysis. AlexaFluor 568 $C_5$ Maleimide (Thermo) was added to DRP1 G401S to reach a ~12-fold molar excess of fluorophore:DRP1 (manufacturer recommends between a 10- to 20-molar excess of fluorophore). This mixture was incubated at 25°C, shaking gently at 200 rpm for 2 hours, as per the manufacturer's specifications. During incubation, a Sephadex G-25 M PD-10 desalting column was pre-equilibrated with 10 column volumes of 0.45-μm filtered dialysis buffer. After the sample was incubated for the allotted time, it was added directly to the equilibrated PD-10 column with additional dialysis buffer to bring the volume to ~1 mL. Fractions were collected in 1 mL increments, eluting with 1 mL of dialysis buffer each time. Fractions believed to contain labeled protein were concentrated in 10 kDa MW cut-off microcentrifuge columns (Amicon) spinning at 14,000 xg at 4°C in 10-minute intervals, until ~200 μL of sample remained. Protein concentration and degree of labeling were determined with equations 1 and 2 using measurements obtained from a NanoDrop One$^{C}$ Microvolume UV-Vis Spectrophotometer. DRP1 G401S (9 cysteines) had a degree of labeling of ~1.66 dye molecules per protein molecule. Labeled protein was aliquoted into single-use 20 μL samples were frozen at -80°C until use.

**GST AlexaFluor 488 $C_5$ Maleimide Labeling**

Purified and cleaved GST (plasmid pGEX-6P-1) was dialyzed overnight into 20 mM HEPES pH 7.4, 150 mM KCl at 4°C. Protein was removed from dialysis and concentrated using a 10 kDa MW cut-off concentrating column (Millipore) centrifuged at 3,300 xg at 4°C in 10-minute intervals to a final concentration of 30 μM . AlexaFluor 488 $C_5$ Maleimide (Thermo) was added to GST to reach a ~15-fold molar excess of fluorophore:GST (manufacturer recommends between a 10- to 20-molar excess of fluorophore). This mixture was incubated at 25°C, shaking gently at 200 rpm for 3.5 hours, as per the manufacturer's specifications. During incubation, a Sephadex G-25 M PD-10 desalting column was pre-equilibrated with 10 column volumes of 0.45-μm filtered dialysis buffer (20 mM HEPES pH 7.4, 150 mM KCl). After the sample was incubated for the allotted time, it was added directly to the equilibrated PD-10 column with additional dialysis buffer to bring the volume to ~1 mL. Fractions were collected in 1 mL increments, eluting with 1 mL of dialysis buffer each time. Fractions believed to contain labeled protein were concentrated in 10 kDa MW cut-off microcentrifuge columns (Amicon) spinning at 4,000 xg at 4°C in 10-minute intervals, until ~200 μL of sample remained. Protein concentration and degree of labeling were determined with equations 1 and 2 using measurements obtained from a NanoDrop One$^{C}$ Microvolume UV-Vis Spectrophotometer. GST (4 cysteines) had a degree of labeling of ~2.90 dye molecules per protein molecule. Labeled protein was aliquoted into single-use 20 μL samples were frozen at -80°C until use.

$$Degree\ of\ labeling = \frac{(A_{max} \times \varepsilon_{280})}{\big(A_{280} - (A_{max} \times CF)\big)} \times \varepsilon_{max} \qquad \text{Equation 1}$$

$$Corrected\ concentration = \frac{(A_{280} - A_{max} \times CF)}{\varepsilon_{280}} \times MW\ protein \qquad \text{Equation 2}$$

**YFP-DRP1 FRAP Microscopy** (continued from Main Text)

FIS1 -/- HCT116 cells were grown in Dulbecco's Modified Eagle Medium:Nutrient Mixture F-12, HEPES (Gibco; 11330-032) supplemented with 10% (v/v) FBS for 24 hours after being plated onto #1.5 high-performance cover glass dishes with 20 mm microwell (Cellvis; D35-20-1.5H) at an initial seeding density of ~400,000 cells per dish. The cells were then transfected with 0.25 μg of a YFP-DRP1-iso3 fusion construct and 1.0 μg of pcDNA, using Avalanche-Omni transfection reagent in Opti-MEM (Gibco; 31985070), for ~16 hours (or overnight) and following manufacturer recommendations. Afterwards, the transfection media was removed, and Dulbecco's Modified Eagle Medium:Nutrient Mixture F-12, HEPES, no phenol red (Gibco; 11039-021) was resupplied.

**YFP-DRP1 Immunofluorescence Microscopy** (continued from Main Text)

Retinal pigmented epithelial (RPE) cells were grown in Dulbecco's Modified Eagle Medium:Nutrient Mixture F-12, HEPES (Gibco; 11330-032) supplemented with 10% (v/v) FBS for 24 hours after being plated onto either a #1.5 high-performance cover glass 24-well plate (Cellvis; P24-1.5H-N) at 37,500 cells/well, or onto a #1.5 high-performance cover glass 6-well plate (Cellvis; P06-1.5H-N) at 150,000 cells/well. The cells were then transfected with 0.20 µg of a YFP-DRP1-iso3 fusion construct (or isoforms 1, 2, or 4 where noted) and 1.05 µg of pcDNA, using Avalanche-Omni transfection reagent in Opti-MEM (Gibco; 31985070), for ~16 hours (or overnight). Afterwards, the transfection media was removed, and the nutrient media was resupplied. 1,6-hexanediol (Sigma-Aldrich; 88571-100ML-F) or 1,2,3-hexanetriol (Sigma-Aldrich; 52895-1G) were diluted in media to achieve the desired 5.0% (v/v) solutions. These solutions were then allowed to equilibrate within the incubator for at least 15 minutes before treatment. Media was aspirated from all wells, proceeding one column at a time. Untreated wells received fresh media, and treated wells received the appropriate 1,6-hexanediol:media or 1,2,3-hexanteiol:media solution for 5 or 20 minutes. The cells were returned to the incubator for the duration of treatment. The cells were protected from light throughout all incubation steps after this point. Following treatment, media and treatment solutions were aspirated from all wells, proceeding one column at a time. Cells were fixed in 4% paraformaldehyde (Thermo; 043368-9M) in 1X PBS (Gibco; 70011-044) for 25 minutes, shaking gently at room temperature. Next, cells were permeabilized in 0.15% (v/v) Triton X-100 (Sigma-Aldrich; X100-1L) in 1X PBS for 15 minutes, shaking gently at room temperature, then blocked in 3% (w/v) BSA (Sigma-Aldrich; A7030-100G) and 0.3% (v/v) Triton X-100 in 1X PBS for 1 hour, shaking gently at room temperature. Primary antibody, rabbit anti-TOM20 (Santa Cruz; sc-11415), was diluted 1:100 in blocking buffer and added to the cells overnight, with gentle rocking at 4°C. Afterward, cells were washed with 1X PBS 3 times for 5 minutes each before receiving the appropriate conjugated secondary antibody, goat anti-rabbit AlexaFluor 568 (Life Technologies; A-11011), diluted 1:500 in blocking buffer for 1 hour, rocking gently at room temperature. Subsequently, cells were washed again with 1X PBS 3 times for 5 minutes each, then received DAPI solution (Thermo; 62248) diluted 1:1000 in blocking buffer for 10 minutes, rocking gently at room temperature. Cells were washed in 1X PBS 3 times for 5 minutes each, then stored in 1X PBS containing 0.2% (w/v) sodium azide (Thermo; S2002-500G), sealed tightly in parafilm, and stored at 4°C until they were imaged.

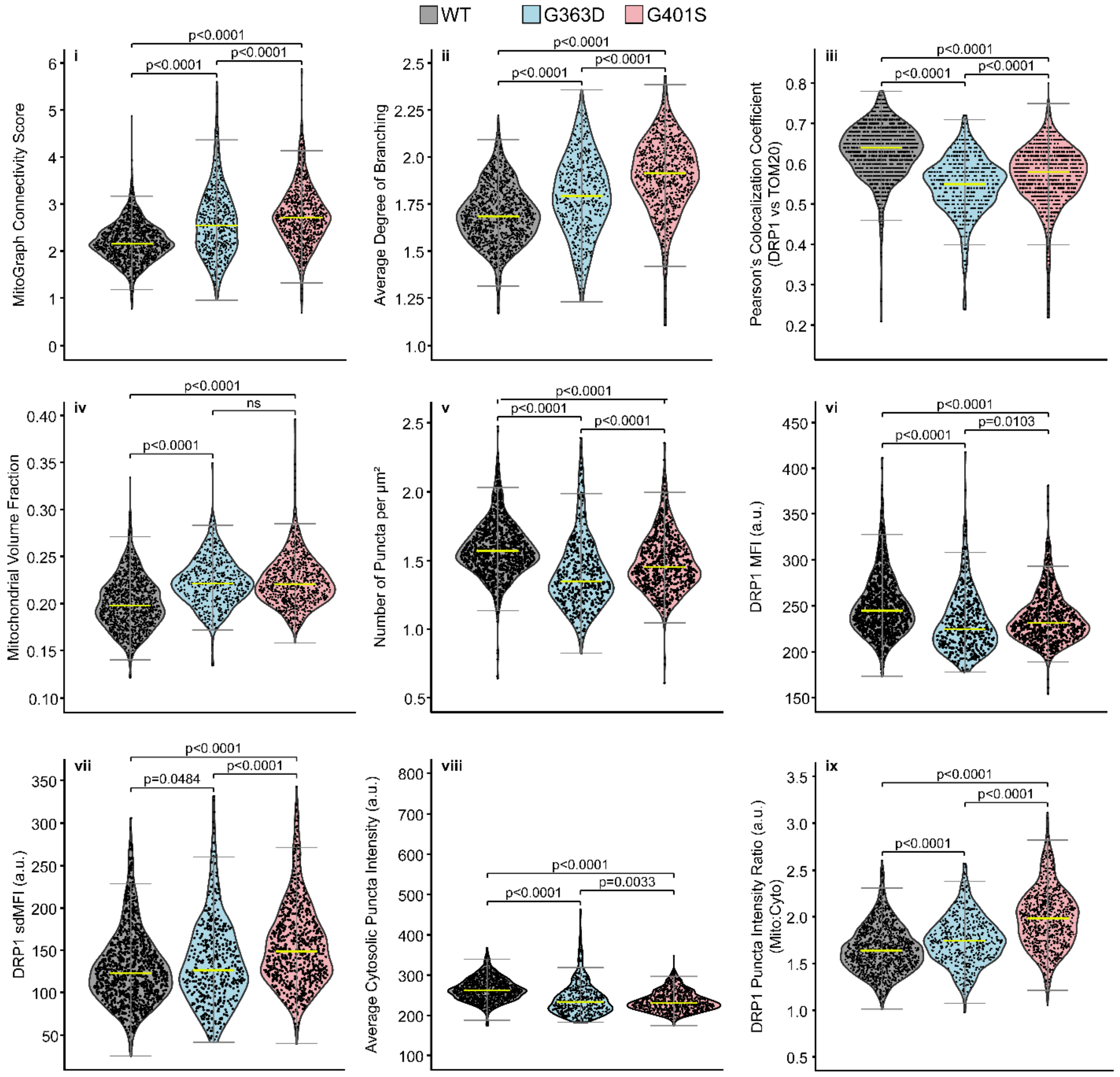


**Supplemental Figure 1.** Additional quantitative metrics collected from cell images of untreated human skin fibroblasts for DRP1 WT (gray), G363D (light blue), and G401S (light red). MitoGraph Connectivity Score (i), average degree of branching (ii), and mitochondrial volume fraction were obtained via MitoGraph software (see main text methods for more information). The MCS is a combinatorial metric that describes the overall complexity of mitochondria within a cell, the value of which increases with more interconnected mitochondrial networks. The degree of branching reported the number of branch points as mitochondrial networks become more interconnected. Pearson's Colocalization coefficients (iii) were calculated using the "coloc2" function in FIJI. Mitochondrial volume fraction (iv) represents the amount of three-dimensional space that mitochondria occupy. Number of puncta per square micron (v) was calculated using the area of the two-dimensional ROI of a given cell. MFI (vi) and sdMFI (vii) were determined for each cell, using the native FIJI "Measure" function, and report on the global distribution of immunofluorescent DRP1 signal. Cytosolic DRP1 puncta intensity (viii) and the ratio between mitochondrial and cytosolic puncta intensities (ix) were collected using the 3DSuite plugin (see main text methods for more information). All statistical tests were performed using one-way ANOVA with Tukey's post-hoc multiple comparisons test, and p-values are indicated for each comparison (ns = not significant). Yellow bars represent the population median. Error bars are ± 2SD.

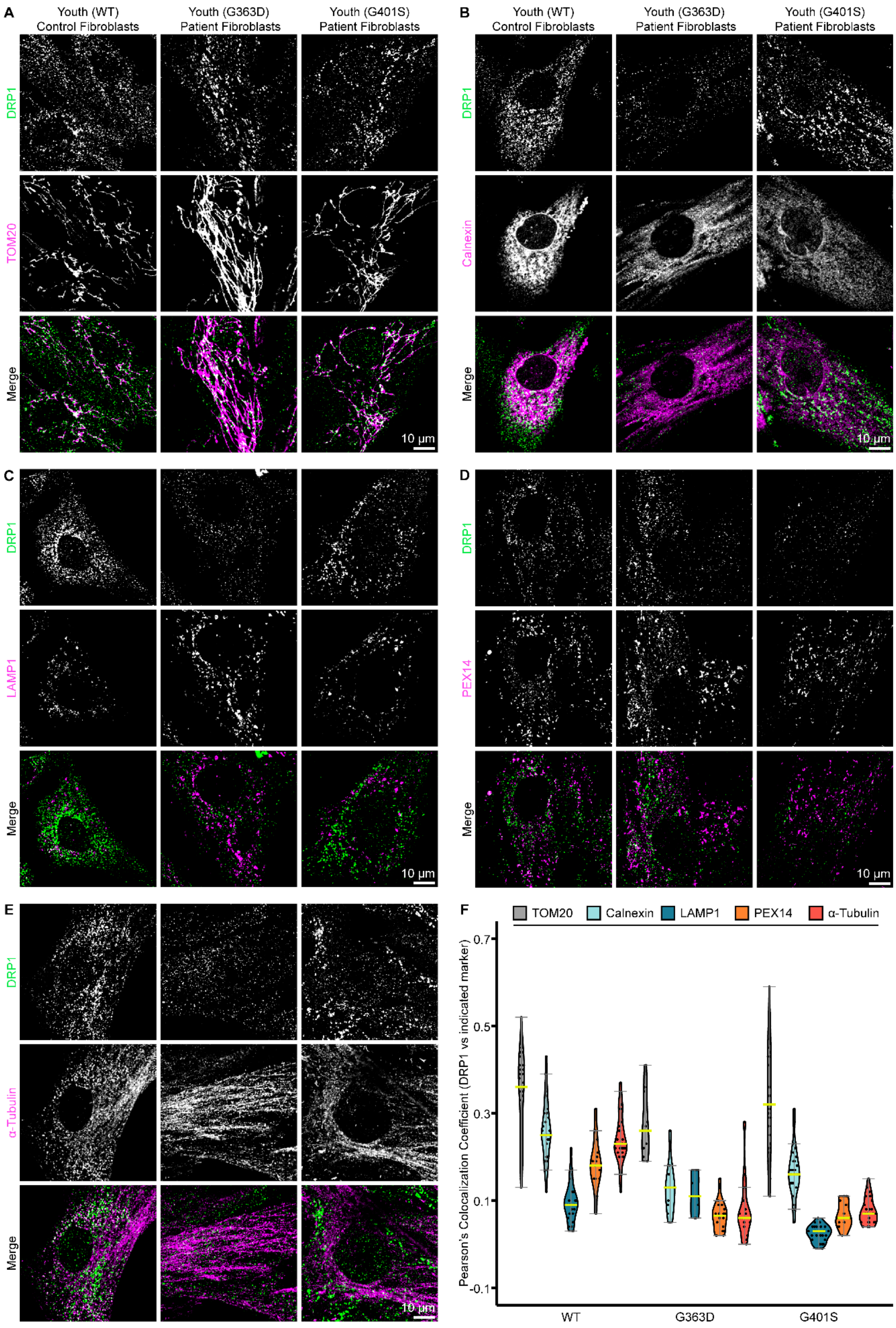
A
Youth (WT) Control Fibroblasts
Youth (G363D) Patient Fibroblasts
Youth (G401S) Patient Fibroblasts
DRP1
TOM20
Merge
10 μm
B
Youth (WT) Control Fibroblasts
Youth (G363D) Patient Fibroblasts
Youth (G401S) Patient Fibroblasts
DRP1
Calnexin
Merge
10 μm
C
DRP1
LAMP1
Merge
10 μm
D
DRP1
PEX14
Merge
10 μm
E
DRP1
α-Tubulin
Merge
10 μm
F
TOM20
Calnexin
LAMP1
PEX14
α-Tubulin
Pearson's Colocalization Coefficient (DRP1 vs indicated marker)
0.7
0.5
0.3
0.1
-0.1
WT
G363D
G401S

**Supplemental Figure 2.** EMPF1 patient fibroblasts and associated age-matched control cells were cultured and prepared for immunofluorescence identically as described in **EMPF1 Patient Fibroblast Culture, Treatment, and Immunolabeling** supplemental materials and methods, with the following exceptions: no samples received 1,6-HD treatment, and cells were immunolabeled with primary antibodies other than rabbit anti-TOM20 where indicated (secondary antibody labeling remained the same since all primary antibodies were of rabbit origin).

Images were captured on a Nikon A1R laser scanning confocal microscope equipped with an A1-DUG Hybrid GaAsP/PMT detector. The microscope was controlled using NIS-Elements Confocal & Enhanced Resolution software package (versions 5.0 and newer), and images were captured on a 60X, 1.40 NA oil objective. The following acquisition settings were utilized for all images: DAPI fluorescence (not shown) was stimulated via 405 nm laser at 10% power, 0 offset, and 110 gain; DRP1 immunofluorescence was stimulated via 488 nm laser at 20% power, -127 offset, and 110 gain; DIC images (not shown) were captured using the transmitted detector via the 488 nm laser, with 0 offset and 80 gain; images were captured at room temperature after a 6.2 second dwell time with 1X averaging through a 12.7 μm pinhole. Alternate antibodies and image acquisition settings are listed per image panel. DRP1 is pseudocolored as green in all images. TOM20, Calnexin, LAMP1, Pex14, and alpha-Tubulin are pseudocolored as magenta where indicated.

Age-matched control cells and EMPF1 patient fibroblasts immunolabeled with mouse anti-DRP1 and one of **(A)** rabbit amti-Tom20 (acquisition settings: 561 nm laser at 10% power, -127 offset, 60 gain), **(B)** rabbit anti-Calnexin (Abcam; ab22595) (acquisition settings: 561 nm laser at 10% power, -127 offset, 80 gain), **(C)** rabbit anti-LAMP1 (Cell Signaling; 9091) (acquisition settings: 561 nm laser at 15% power, -127 offset, 100 gain), **(D)** rabbit anti-PEX14 (Abcam; ab183885) (acquisition settings: 561 nm laser at 15% power, -127 offset, 110 gain), or **(E)** rabbit anti-**α**-Tubulin (Cell Signaling; 2144) (acquisition settings: 561 nm laser at 25% power, -127 offset, 150 gain). DRP1 LUTs are identical across all panel sets. All other LUTs are identical within their panel set. Scale bar = 10 μm.

**(F)** Pearson's Colocalization coefficients were calculated using the "coloc2" function in FIJI (described in methods) for DRP1 versus TOM20 (gray), Calnexin (light blue), LAMP1 (teal), PEX14 (orange), and α-Tubulin (red). Yellow bars represent the population median. Error bars are ± 2SD.

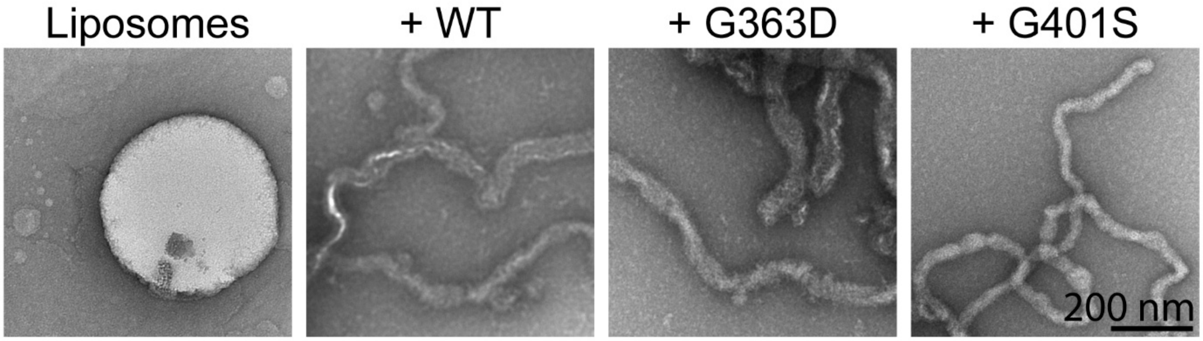


**Supplemental Figure 3.** Electron Microscopy images of 150 μM CL-containing liposomes (40%, 35%, and 25% molar ratios of DOPC, DOPE, and CL, respectively) either alone (leftmost) or with one of 4 μM DRP1 WT (center-left), G363D (center-right), or G401S (rightmost) in Buffer B (20 mM HEPES, pH 7.4; 150 mM KCl; 1 mM DTT). Scale bar = 200 nm.

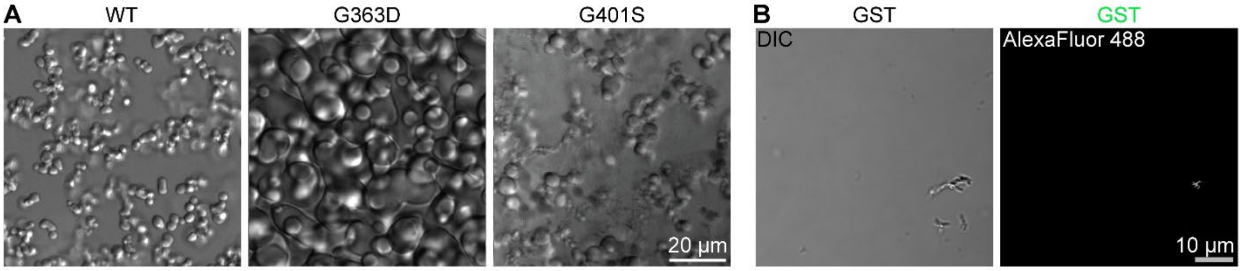


**Supplemental Figure 4. (A)** DIC images of 25 μM DRP1 constructs with 20% (w/v) Ficoll PM 70 (Sigma; F2878-50G) in DRP1 phase separation buffer (20 mM HEPES, pH 7.4; 150 mM KCl; 2 mM $MgCl_2$; 1 mM DTT). These images are representative of several experiments completed at various concentrations of DRP1 and crowder. Scale bar = 20 μm. **(B)** DIC and fluorescence images of 20 μM GST (5% labeled with AlexaFluor 488) in GST phase separation buffer (20 mM HEPES, pH 7.4; 150 mM KCl; 1 mM DTT) containing 5% (w/v) PEG-8K. Scale bar = 10 μm.

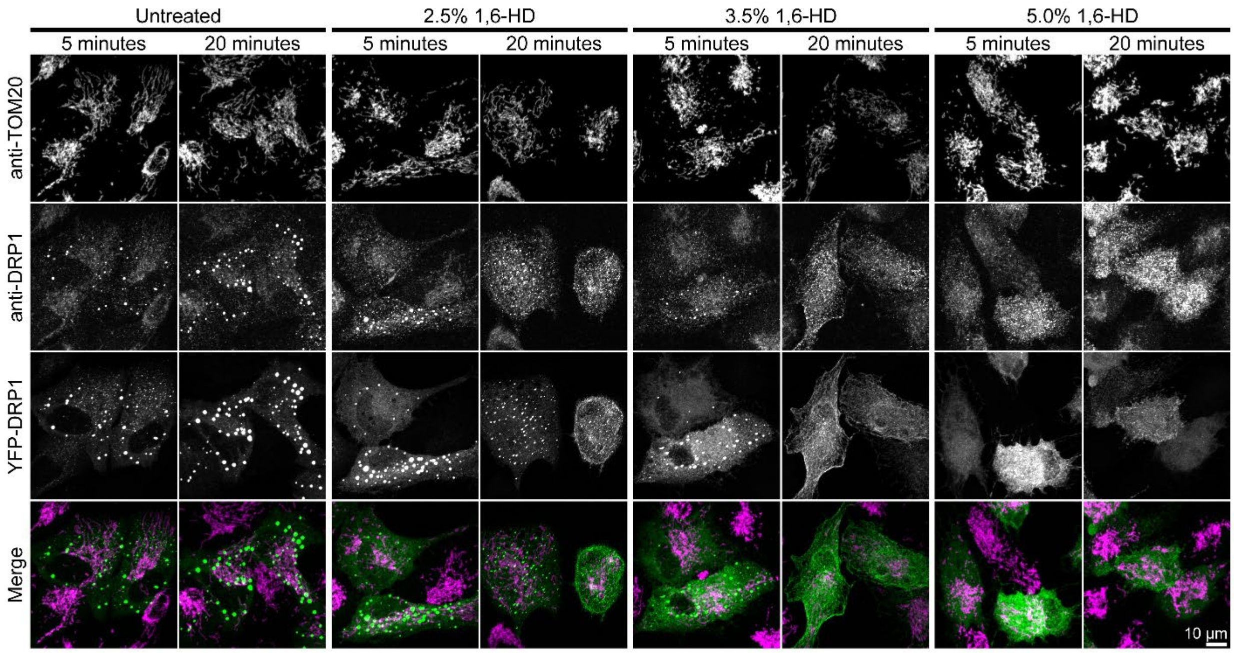


**Supplemental Figure 5.** RPE cells were grown in Dulbecco's Modified Eagle Medium:Nutrient Mixture F-12, HEPES (Gibco) supplemented with 10% (v/v) FBS for 24 hours after being plated onto #1.5 high-performance cover glass 96-well plates (Cellvis) at an initial seeding density of ~37,500 cells per well. The cells were then transfected with 1.25 µg of a YFP-DRP1-iso3 fusion construct, using Avalanche-Omni transfection reagent in Opti-MEM (Gibco), for ~16 hours (or overnight). Afterwards, the transfection media was removed, and the nutrient media was resupplied. The desired 1,6-HD concentrations (v/v) were achieved by dilution in nutrient media before their addition to the cells. Nutrient media was aspirated from all wells, proceeding one column at a time. Untreated wells received fresh media, and treated wells received the appropriate 1,6-hexanediol:media solution for the allotted incubation time. The cells were returned to the incubator for the duration of treatment.

Following treatment, media and treatment solutions were aspirated from all wells, proceeding one column at a time. Cells were fixed in 4% paraformaldehyde (Thermo) in 1X PBS (Gibco) for 25 minutes, shaking gently at room temperature. Next, cells were permeabilized in 0.15% (v/v) Triton X-100 (Sigma-Aldrich) in 1X PBS for 15 minutes, shaking gently at room temperature, then blocked in 3% (w/v) BSA (Sigma-Aldrich) and 0.3% (v/v) Triton X-100 in 1X PBS for 1 hour, shaking gently at room temperature. Primary antibody, mouse anti-Drp1 (BD Biosciences; 611113) and rabbit anti-Tom20 (Santa Cruz; sc-11415), were diluted 1:100 in blocking buffer and added to the cells overnight, with gentle rocking at 4°C. Afterward, cells were washed with 1X PBS 3 times for 5 minutes each before receiving the appropriate conjugated secondary antibody, either goat anti-mouse AlexaFluor 568 (Life Technologies; A-11004) or goat anti-rabbit AlexaFluor 647 (Invitrogen; A-32733), diluted 1:500 in blocking buffer for 1 hour, rocking gently at room temperature. Subsequently, cells were washed again with 1X PBS 3 times for 5 minutes each, then received DAPI solution (Thermo; 62248) diluted 1:1000 in blocking buffer for 10 minutes, rocking gently at room temperature. Cells were washed in 1X PBS 3 times for 5 minutes each, then stored in 1X PBS containing 0.2% (w/v) sodium azide (Thermo; S2002-500G), sealed tightly in parafilm, and stored at 4°C until they were imaged.

Images were captured on a Nikon Eclipse Ti microscope base equipped with a Yokogawa CSU-W1 spinning disk confocal scanner unit (50 µm pinhole), 40X 0.90 NA objective, and Hamamatsu ORCAFlash4.0 V3 sCMOS camera. Fluorescent protein and fluorophore activation were stimulated using 488, 561, and 647 nm lasers, each at 50% power, with images being captured after a 200-millisecond exposure with 2X averaging enabled. Scale bar = 10 µm.

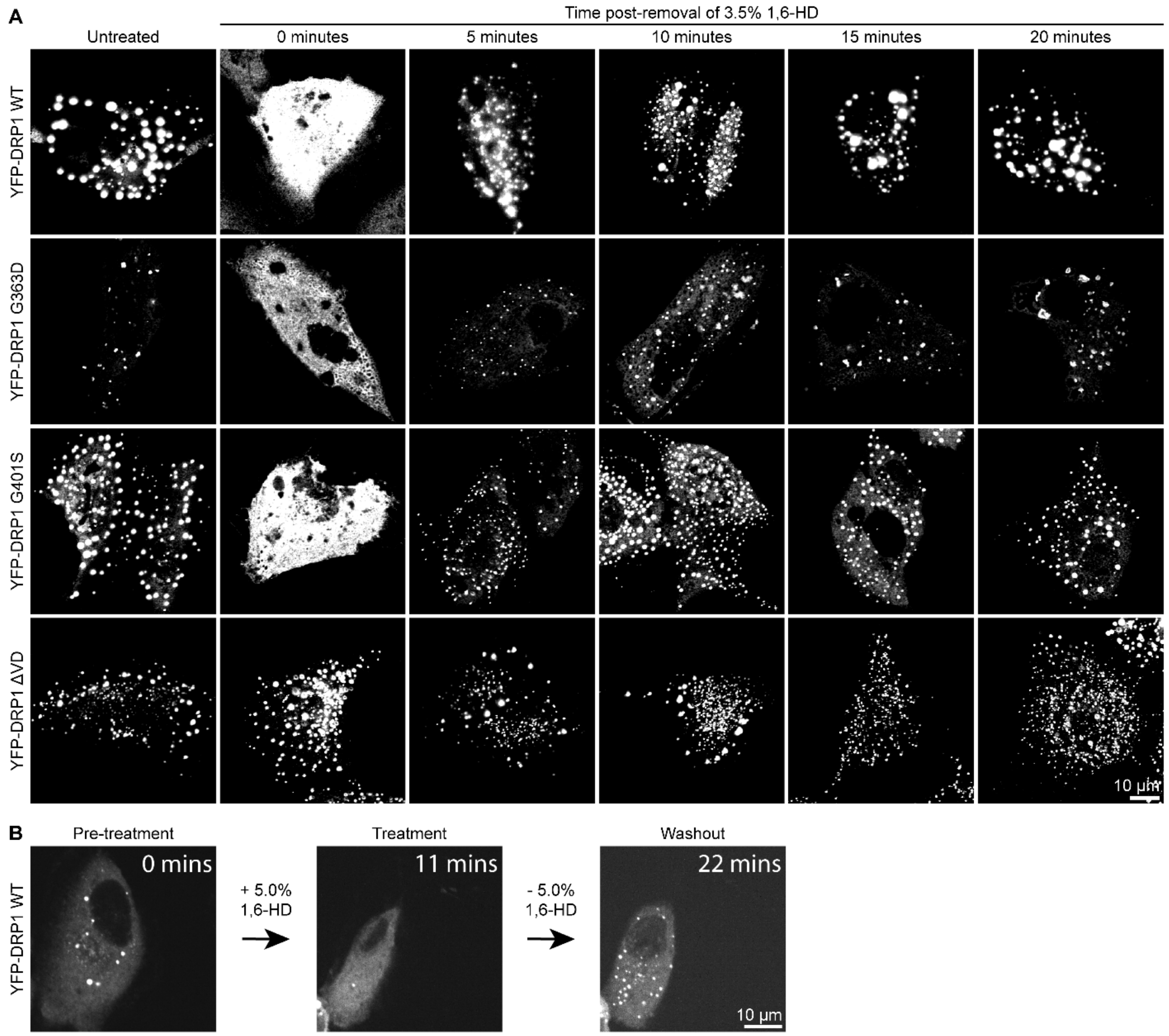


**Supplemental Figure 6. (A)** RPE cells were grown in Dulbecco's Modified Eagle Medium:Nutrient Mixture F-12, HEPES (Gibco; 11330-032) supplemented with 10% (v/v) FBS for 24 hours after being plated onto a #1.5 high-performance cover glass 24-well plate (Cellvis; P24-1.5H-N) at 37,500 cells/well. The cells were then transfected with 1.25 µg of the specified YFP-DRP1 (isoform 1) fusion construct and 1.25 µg of empty vector pcDNA, using Avalanche-Omni transfection reagent (EZ Biosystems; EZT-OMNI-1) in Opti-MEM (Gibco; 31985070), for ~16 hours (or overnight) and following manufacturer recommendations. Afterwards, the transfection media was removed, and the nutrient media was resupplied. 1,6-hexanediol (Sigma-Aldrich; 88571-100ML-F) was diluted in media to achieve the desired 3.5% or 5.0% (v/v) solution. This solution was then allowed to equilibrate within the incubator for at least 15 minutes before treatment. Media was aspirated from all wells, proceeding one column at a time. Untreated wells received fresh media, and treated wells received the 1,6-hexanediol:media solution for 5 minutes. The cells were returned to the incubator for the duration of treatment. Afterwards, the 1,6-hexanediol:media solution was removed and replaced with fresh media for the specified duration.

Cells were then fixed in 4% paraformaldehyde (Thermo; 043368-9M) in 1X PBS (Gibco; 70011-044) for 25 minutes, shaking gently at room temperature. Next, cells were permeabilized in 0.15% (v/v) Triton X-100 (Sigma-Aldrich; X100-1L) in 1X PBS for 15 minutes, shaking gently at room temperature, then blocked in 3% (w/v) BSA (Sigma-Aldrich; A7030-100G) and 0.3% (v/v) Triton X-100 in 1X PBS for 1 hour, shaking gently at room temperature. Primary antibody, rabbit anti-TOM20 (Santa Cruz; sc-11415), was diluted 1:100 in blocking buffer and added to the cells overnight, with gentle rocking at 4°C. Afterward, cells were washed with 1X PBS 3 times for 5 minutes each before receiving the appropriate conjugated secondary antibody, goat anti-rabbit AlexaFluor 568 (Life Technologies; A-11011), diluted 1:500 in blocking buffer for 1 hour, rocking gently at room temperature. Subsequently, cells were washed again with 1X PBS 3 times for 5 minutes each, then received DAPI solution (Thermo; 62248) diluted 1:1000 in blocking buffer for 10 minutes, rocking gently at room temperature. Cells were washed in 1X PBS 3 times for 5 minutes each, then stored in 1X PBS containing 0.2% (w/v) sodium azide (Thermo; S2002-500G), sealed tightly in parafilm, and stored at 4°C until they were imaged.

Images were captured on a Nikon A1R laser scanning confocal microscope equipped with an A1-DUG Hybrid GaAsP/PMT detector. The microscope was controlled using NIS-Elements Confocal & Enhanced Resolution software package (versions 5.0 and newer), and images were captured on a 60X, 1.40 NA oil objective. DAPI fluorescence was stimulated via 405 nm laser at 10% power, 0 offset, and 110 gain. YFP-DRP1 fluorescence was stimulated via 488 nm laser at 15% power, -127 offset, and 70 gain. TOM20 immunofluorescence was stimulated via 561 nm laser at 15% power, -127 offset, and 70 gain. DIC images were captured using the transmitted detector via the 488 nm laser, with 0 offset and 80 gain. All images were captured at room temperature after a 6.2 second dwell time with 1X averaging through a 12.7 μm pinhole. LUTs are identical across all images. Scale bar = 10 μm.

**(B)** RPE cells were plated and transfected similarly as above, except the cells were plated onto #1.5 high-performance cover glass 4-chamber 35 mm dishes with 20 mm microwell (Cellvis; D35C4-20-1.5-N) at an initial seeding density of ~37,500 cells per chamber, and the cells were transfected with 0.3125 μg of YFP-DRP1-iso3 fusion construct (and 0.3125 μg of pcDNA) instead.

Confocal fluorescence microscopy images were captured on a Nikon Eclipse Ti microscope base equipped with a Yokogawa CSU-W1 spinning disk confocal scanner unit (50 μm pinhole), 40X 0.90 NA objective, and Hamamatsu ORCAFlash4.0 V3 sCMOS camera. The microscope environmental chamber was pre-equilibrated to 37°C, 5% $CO_2$, and ~90% humidity prior to staging of the dish. Additionally, the nutrient media was aspirated from each chamber and replaced with Dulbecco's Modified Eagle Medium:Nutrient Mixture F-12, HEPES, no phenol red (Gibco; 11039-021).

Fluorescent protein activation was stimulated using a 488 nm laser at 50% power, with images being captured after a 200-millisecond exposure. A time series was established such that an image was acquired once per minute. At the 10-minute mark, the media was carefully aspirated from the chamber of the dish and was replaced with media containing 5% (v/v) 1,6-HD. Imaging continued for another 10 minutes. Afterwards, the media was once again aspirated from the chamber of the dish, this time being replaced with fresh media (not containing 1,6-HD). Imaging continued for an additional 10 minutes. Images corresponding to the first minute after startup and media swaps were selected to highlight the speed at which DRP1 puncta were influenced by 1,6-HD. Scale bar = 10 μm.

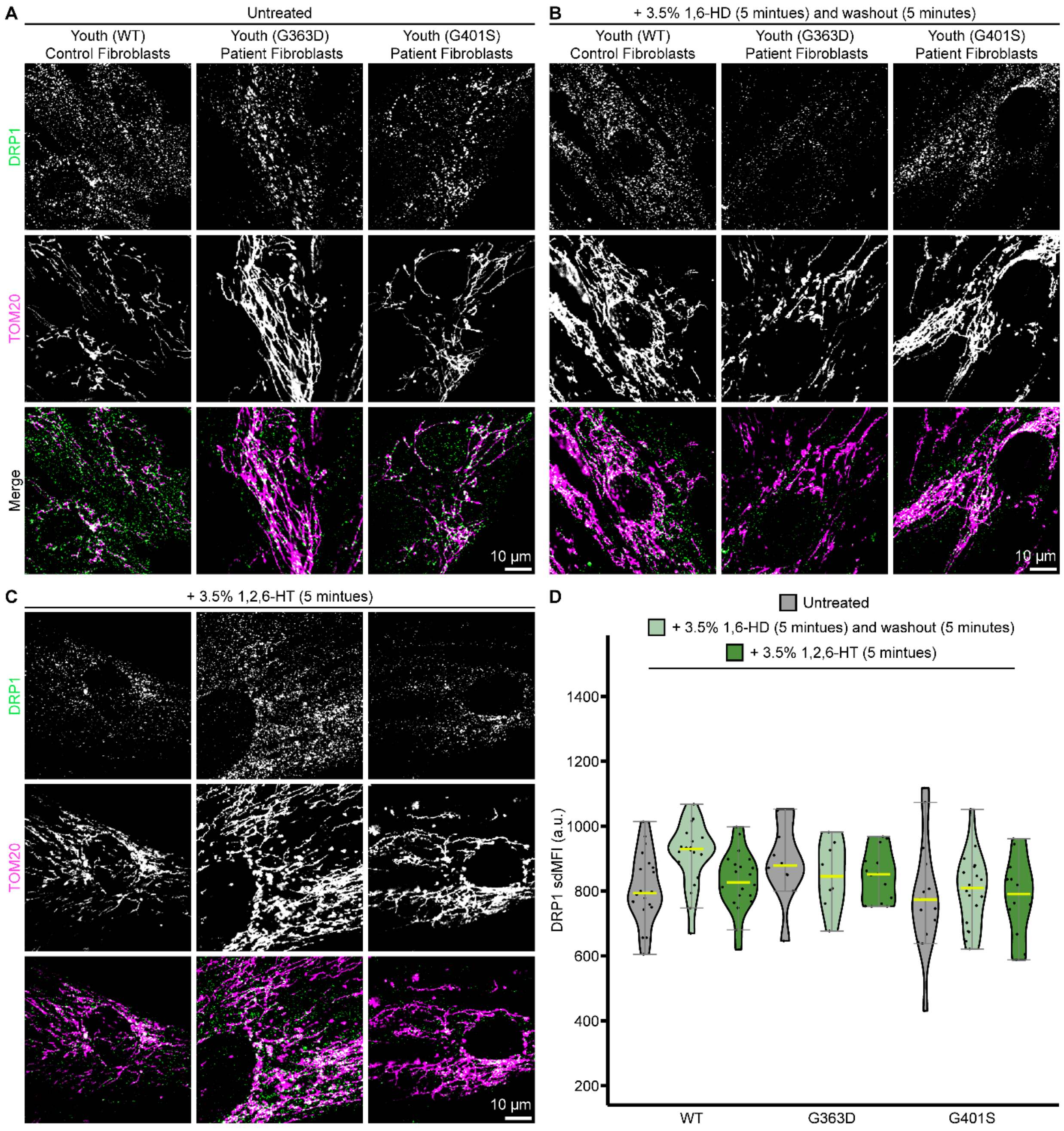


**Supplemental Figure 7.** EMPF1 patient fibroblasts and associated age-matched control cells were cultured and prepared for immunofluorescence identically as described in **EMPF1 Patient Fibroblast Culture, Treatment, and Immunolabeling** supplemental materials and methods, with exceptions in the treatment regimen as indicated in **(B)** and **(C)**.

Images were captured on a Nikon A1R laser scanning confocal microscope equipped with an A1-DUG Hybrid GaAsP/PMT detector. The microscope was controlled using NIS-Elements Confocal & Enhanced Resolution software package (versions 5.0 and newer), and images were captured on a 60X, 1.40 NA oil objective. The following acquisition settings were utilized for all images: DAPI fluorescence (not shown) was stimulated via 405 nm laser at 10% power, 0 offset, and 110 gain; DRP1 immunofluorescence was stimulated via 488 nm laser at 20% power, -127 offset, and 110 gain; TOM20 immunofluorescence was stimulated via 561 nm laser at 10% power, -127 offset, and 60 gain; DIC images (not shown) were captured using the transmitted detector via the 488 nm laser, with 0 offset and 80 gain; images were captured at room temperature after a 6.2 second dwell time with 1X averaging through a 12.7 µm pinhole.

Cell types of DRP1 WT (left), DRP1 G363D (center), and DRP1 G401S (right) are organized by column for each panel set. Samples were either untreated **(A)**, treated with 3.5% 1,6-hexanediol for 5 minutes followed by washout and equilibration with nutrient media for an additional 5 minutes **(B)**, or treated with 3.5% 1,2,6-hexanetriol for 5 minutes **(C)**. DRP1 is pseudocolored as green, and TOM20 is pseudocolored as magenta in all images. LUTs are identical across all panel sets. Scale bar = 10 μm.

**(D) The** sdMFI was calculated for each cell, using the native FIJI “Measure” function, and reported on the global distribution of immunofluorescent DRP1 signal for cells that were either untreated (gray), treated with 3.5% 1,6-hexanediol for 5 minutes followed by washout and equilibration with nutrient media for an additional 5 minutes (light green), or treated with 3.5% 1,2,6-hexanetriol for 5 minutes (dark green). Cell types are indicated on the X-axis and were grouped as a triplicate set. Yellow bars represent the population median. Error bars are ± 2SD.

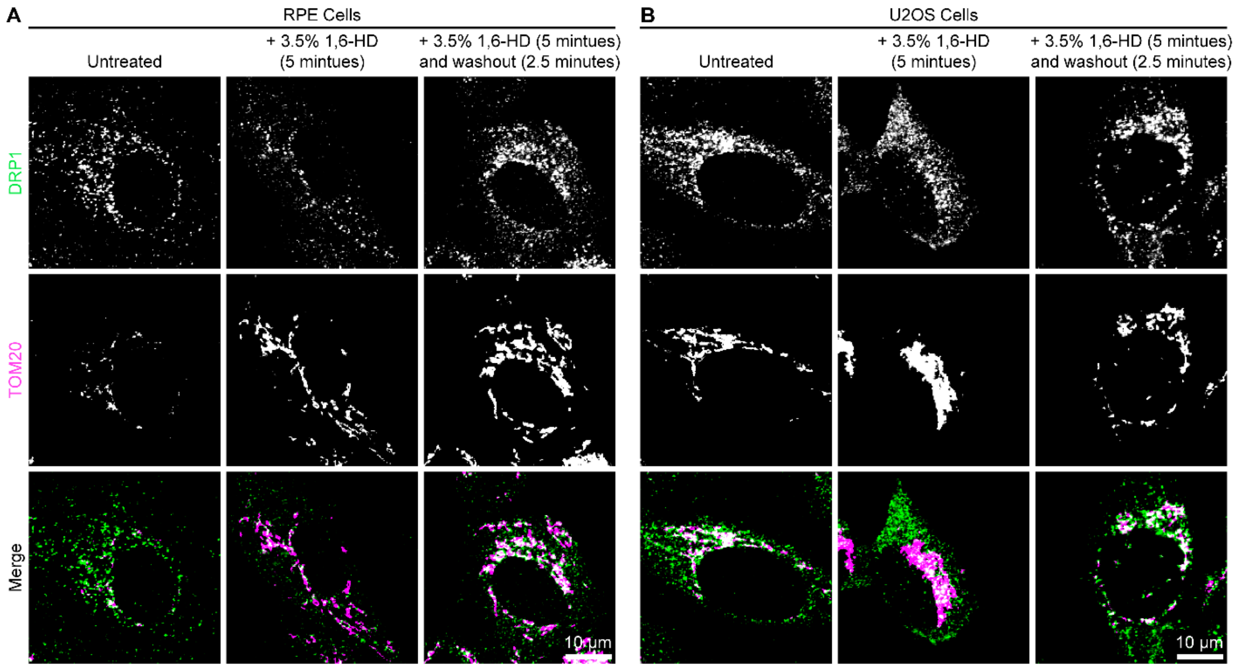


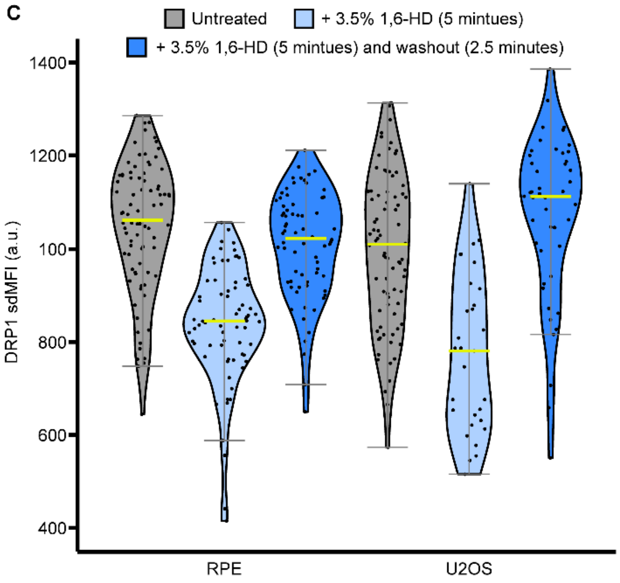


**Supplemental Figure 8.** RPE cells were grown in Dulbecco's Modified Eagle Medium:Nutrient Mixture F-12, HEPES (Gibco; 11330-032) supplemented with 10% (v/v) FBS for 48 hours after being plated onto a #1.5 high-performance cover glass 24-well plate (Cellvis; P24-1.5H-N) at 37,500 cells/well. U2OS cells were grown identically as above, but were plated at a density of 25,000 cells/well instead.

1,6-hexanediol (Sigma-Aldrich; 88571-100ML-F) was diluted in media to achieve the desired 3.5% (v/v) solution. This solution was then allowed to equilibrate within the incubator for at least 15 minutes before treatment. Media was aspirated from all wells, proceeding one column at a time. Untreated wells received fresh media, and treated wells received the 1,6-hexanediol:media solution for 5 minutes. The cells were returned to the incubator for the duration of treatment. Afterwards for washouts, the 1,6-hexanediol:media solution was removed and replaced with fresh media for 2.5 minutes.

Cells were then fixed in 4% paraformaldehyde (Thermo; 043368-9M) in 1X PBS (Gibco; 70011-044) for 25 minutes, shaking gently at room temperature. Next, cells were permeabilized in 0.15% (v/v) Triton X-100 (Sigma-Aldrich; X100-1L) in 1X PBS for 15 minutes, shaking gently at room temperature, then blocked in 3% (w/v) BSA (Sigma-Aldrich; A7030-100G) and 0.3% (v/v) Triton X-100 in 1X PBS for 1 hour, shaking gently at room temperature. Primary antibody, rabbit anti-TOM20 (Santa Cruz; sc-11415), was diluted 1:100 in blocking buffer and added to the cells overnight, with gentle rocking at 4°C. Afterward, cells were washed with 1X PBS 3 times for 5 minutes each before receiving the appropriate conjugated secondary antibody, goat anti-rabbit AlexaFluor 568 (Life Technologies; A-11011), diluted 1:500 in blocking buffer for 1 hour, rocking gently at room temperature. Subsequently, cells were washed again with 1X PBS 3 times for 5 minutes each, then received DAPI solution (Thermo; 62248) diluted 1:1000 in blocking buffer for 10 minutes, rocking gently at room temperature.

(continued from text in figure) Cells were washed in 1X PBS 3 times for 5 minutes each, then stored in 1X PBS containing 0.2% (w/v) sodium azide (Thermo; S2002-500G), sealed tightly in parafilm, and stored at 4°C until they were imaged.

Images were captured on a Nikon A1R laser scanning confocal microscope equipped with an A1-DUG Hybrid GaAsP/PMT detector. The microscope was controlled using NIS-Elements Confocal & Enhanced Resolution software package (versions 5.0 and newer), and images were captured on a 60X, 1.40 NA oil objective. The following acquisition settings were utilized for all images: DAPI fluorescence (not shown) was stimulated via 405 nm laser at 10% power, 0 offset, and 110 gain; DRP1 immunofluorescence was stimulated via 488 nm laser at 20% power, -127 offset, and 110 gain; TOM20 immunofluorescence was stimulated via 561 nm laser at 10% power, -127 offset, and 60 gain; DIC images (not shown) were captured using the transmitted detector via the 488 nm laser, with 0 offset and 80 gain; images were captured at room temperature after a 6.2 second dwell time with 1X averaging through a 12.7 μm pinhole.

RPE **(A)** and U2OS **(B)** cells were either untreated (left column), treated with 3.5% 1,6-hexanediol for 5 minutes (center column), or **treated with** 3.5% 1,6-hexanediol for 5 minutes followed by washout and equilibration with nutrient media for an additional 2.5 minutes (right column) for each panel set. DRP1 is pseudocolored as green, and TOM20 is pseudocolored as magenta in all images. LUTs are identical across all panel sets. Scale bar = 10 μm.

**(C) The** sdMFI was calculated for each cell, using the native FIJI “Measure” function, and reported on the global distribution of immunofluorescent DRP1 signal for untreated (gray), 3.5% 1,6-hexanediol treated for 5 minutes (light blue), 3.5% 1,6-hexanediol treated for 5 minutes followed by washout and equilibration with nutrient media for an additional 2.5 minutes (dark blue). Yellow bars represent the population median. Error bars are ± 2SD.

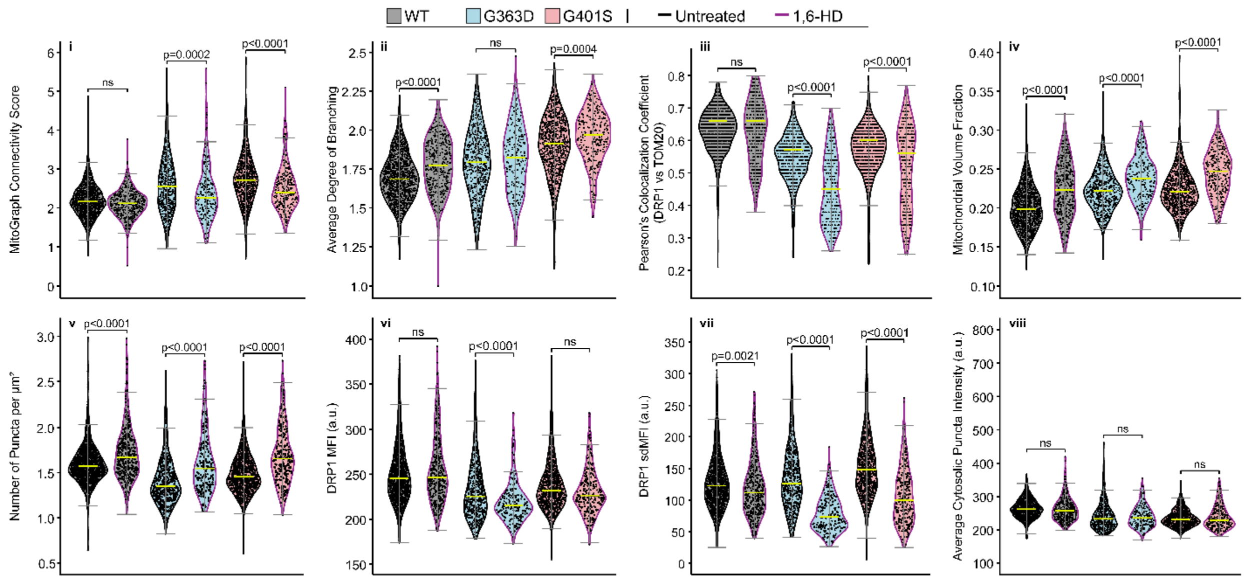


**Supplemental Figure 9.** Additional quantitative metrics collected from cell images of untreated (black outline) and 3.5% (v/v) 1,6-HD treated (magenta outline) human skin fibroblasts for DRP1 WT (gray), G363D (light blue), and G401S (light red). MitoGraph Connectivity Score (i), average degree of branching (ii), and mitochondrial volume fraction were obtained via MitoGraph software (see main text methods for more information). The MCS is a combinatorial metric that describes the overall complexity of mitochondria within a cell, the value of which increases with more interconnected mitochondrial networks. The degree of branching reported the number of branch points as mitochondrial networks become more interconnected. Pearson's Colocalization coefficients (iii) were calculated using the "coloc2" function in FIJI. Mitochondrial volume fraction (iv) represents the amount of three-dimensional space that mitochondria occupy. Total number of DRP1 puncta per square micron (v) was calculated using the area of the two-dimensional ROI of a given cell with the "3DSuite" plugin in FIJI (see main text methods for more information). MFI (vi) and sdMFI (vii) were calculated for each cell, using the native FIJI "Measure" function, and reported on the global distribution of immunofluorescent DRP1 signal. Cytosolic DRP1 puncta intensities (viii) were collected using the 3DSuite plugin (see main text methods for more information). All statistical tests were performed using one-way ANOVA with Tukey's post-hoc multiple comparisons test, and p-values are indicated for each comparison (ns = not significant). Yellow bars represent the population median. Error bars are ± 2SD.

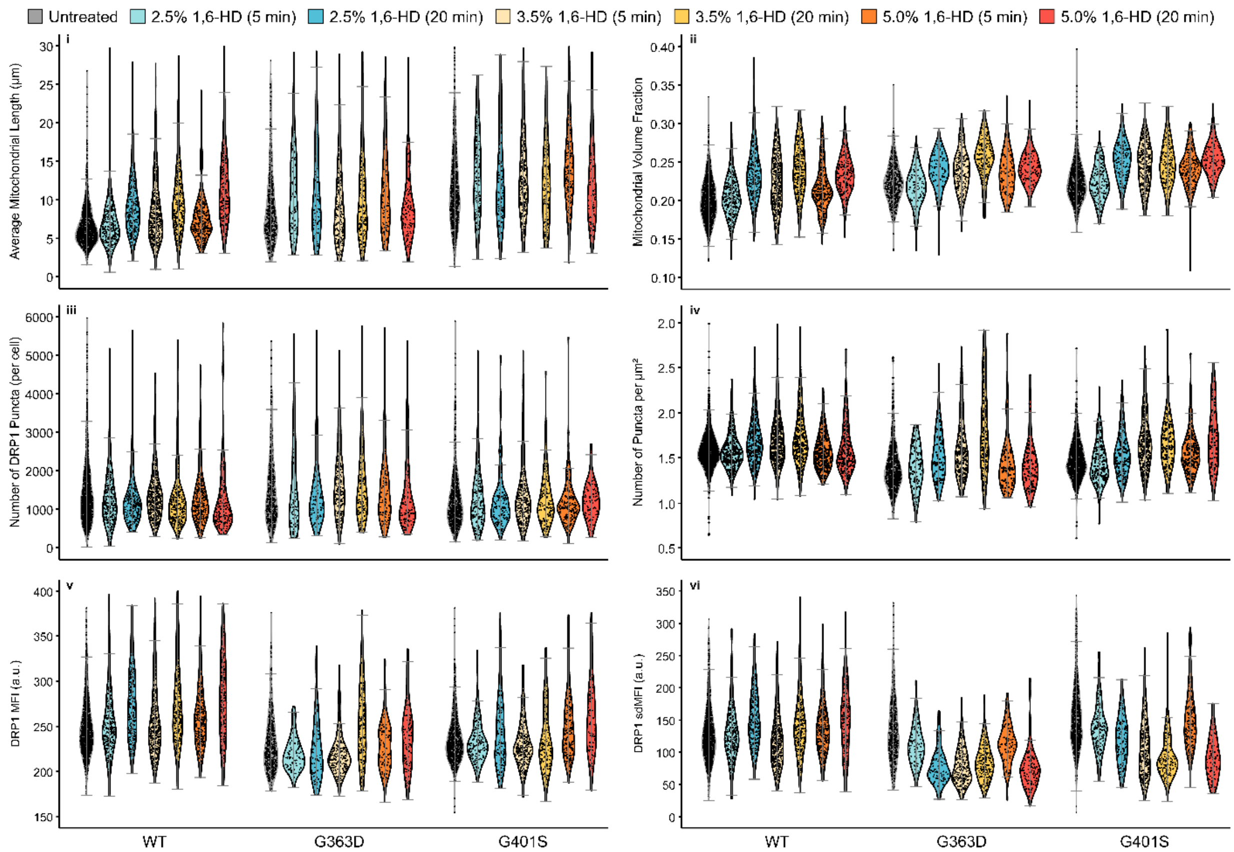


**Supplemental Figure 10.** Additional quantitative metrics collected from cell images of human skin fibroblasts either untreated (gray), 2.5% 1,6-HD for 5 minutes (light blue), 2.5% 1,6-HD for 20 minutes (blue), 3.5% 1,6-HD for 5 minutes (light yellow), 3.5% 1,6-HD for 20 minutes (yellow), 5.0% 1,6-HD for 5 minutes (orange), or 5.0% 1,6-HD for 20 minutes (red) for DRP1 WT (left facet), G363D (middle facet), or G401S cells (right facet). Average mitochondrial length (iv) and mitochondrial volume fraction (ii) were obtained via MitoGraph software (see main text methods for more information). The total number of DRP1 puncta (iii) was collected using the 3DSuite plugin (see main text methods for more information). Number of puncta per square micron (iv) was calculated using the area of the two-dimensional ROI of a given cell. DRP1 MFI (v) and sdMFI (vi) were calculated using the native FIJI “Measure” function. Statistical comparisons were not performed at this scale, but the trends align with those from the main text data (3.5% 1,6-HD for 5 minutes).

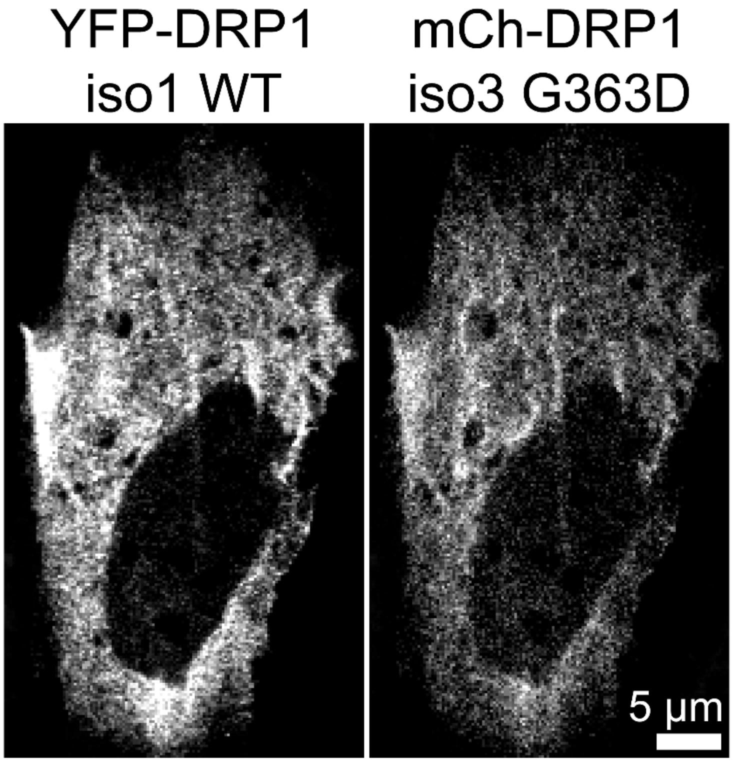


Pearson's Colocalization Coefficient: 0.79
Manders' M1: 0.964
Manders' M2: 0.968

**Supplemental Figure 11.** RPE cells were grown in Dulbecco's Modified Eagle Medium:Nutrient Mixture F-12, HEPES (Gibco) supplemented with 10% (v/v) FBS for 24 hours after being plated onto #1.5 high-performance cover glass 4-chamber 35 mm dishes with 20 mm microwell (Cellvis; D35C4-20-1.5-N) at an initial seeding density of ~37,500 cells per chamber. The cells were then transfected with 0.625 μg of YFP-DRP1-iso1-WT and 0.625 μg of mCherry-DRP1-iso3-G363D fusion constructs with Avalanche-Omni transfection reagent in Opti-MEM (Gibco), for ~16 hours (or overnight) and following manufacturer recommendations. Afterwards, the cells were fixed in 4% paraformaldehyde (Thermo) in 1X PBS (Gibco) for 25 minutes, shaking gently at room temperature.

Images were captured on a Nikon A1R laser scanning confocal microscope equipped with an A1-DUG Hybrid GaAsP/PMT detector. The microscope was controlled using NIS-Elements Confocal & Enhanced Resolution software package (versions 5.0 and newer), and images were captured on a 60X, 1.40 NA oil objective. Fluorescent protein activation was stimulated using 488 and 561 nm lasers, both of which were set to 10% power, 127 offset, and 110 gain. Images were captured at room temperature after a 6.2 second dwell time with 1X averaging through a 12.7 μm pinhole. Scale bar = 5 μm.
